# Ecological drivers and evolutionary consequences of torpor in Andean hummingbirds

**DOI:** 10.1101/2022.12.08.519654

**Authors:** Diana Carolina Revelo Hernández, Justin W. Baldwin, Gustavo A. Londoño

**Author notes:** Address for correspondence: Justin Wheeler Baldwin, Department of Biology, Washington University, St. Louis, MO 63130, USA, CB 8226. Shared first authorship (Diana Carolina Revelo Hernández, Justin W. Baldwin). Author contributions: DCRH and GAL conceived and designed the study. DCRH collected field data. JWB and DCRH performed analyses. JWB collected literature data and wrote the first draft. All authors contributed substantially to revisions. Data accessibility: Data and code are archived on Zenodo under DOI: 10.5281/zenodo.7098176.

## Abstract

Daily torpor allows endotherms to save energy during energetically stressful (e.g., cold) conditions. Although studies on avian torpor have mostly been conducted under laboratory conditions, information on the usage of torpor in the wild is limited to few, predominantly temperate-zone species. We studied torpor under seminatural conditions from 249 individuals from 29 hummingbird species across a 1,920 m elevational gradient in the western Andes of Colombia using cloacal thermistors. Small birds used torpor more often than large birds, but only at low ambient temperatures, where torpor was prolonged. We also found effects of proxy variables for body condition and energy expenditure on the use of torpor, its characteristics, and impacts. Our results suggest that judicious deployment and subtle modifications of torpor, as well as phylogenetic variation in propensity for torpor, have allowed only certain clades of hummingbirds to colonize and persist in the harsh climate of the high Andes.

## Introduction

The tropical Andes reach over 6,700 m above sea level and are the world’s top hotspot of biodiversity (Myers *et al*. 2000). About half of all extant hummingbirds live in the Andes (McGuire *et al*. 2014), and they are found from the warm, moist lowlands to the cold highland páramo. As life at high altitudes demands extreme adaptations, high-altitude hummingbirds endure the thin, hypoxic montane air by altering wing-loading (Altshuler *et al*. 2004a, b; Stiles 2004), efficiently harvesting oxygen with modified hemoglobin (Projecto-Garcia *et al*. 2013), and, occasionally, walking (Vuilleumier 1969). Ascending along tropical elevational gradients, mean ambient temperature (T_a_) decreases and its circadian variation increases, making conditions on tropical mountaintops simultaneously cold and harsh (Janzen 1967). Small-bodied endotherms share high lower limits (T_lc_) of the thermoneutral zone (TNZ; Fristoe *et al*., 2015; Porter and Kearney, 2009), so in cold environments they must expend considerable energy to maintain high body temperatures (T_b_) and thermoregulate (Wolf & Walsberg 2000). However, hummingbirds can alleviate large (and costly) thermal gradients between low T_a_ and high T_b_ by roosting in buffered caves (Pearson 1953) or by deploying torpor during cold nights (e.g. Carpenter, 1974; Pearson, 1950; Wolf *et al*., 2020).

Daily torpor is the facultative reduction in T_b_ to well below normothermy (often near T_a_) during nightly resting periods. Unlike hypothermia, torpor includes an increase in T_b_, that requires birds to produce at least some endogenous heat (Geiser *et al*. 2014); in addition, they can also make use of passive solar warming (Wolf *et al*. 2020). Unlike hibernation, which occurs seasonally, torpor can occur daily (Geiser 2004; Ruf & Geiser 2015). However, hummingbirds calibrate many aspects of torpor in response to ecological conditions such as changes in ambient temperature (Spence & Tingley 2021), weather (Calder & Booser 1973), migration status (Carpenter & Hixon 1988), nutrition (Hainsworth *et al*. 1977; Powers *et al*. 2003), and season (Hiebert 1991). Moreover, the use of torpor appears to be distributed along a continuum (Schleucher 2004; Ruf & Geiser 2015; Shankar *et al*. 2022), and subtle modifications in the frequency of torpor and bout duration are crucial to birds’ ability to save energy (Shankar *et al*. 2020b; Wolf *et al*. 2020). As a consequence, the use of torpor varies across species (Krüger *et al*. 1982; Bech *et al*. 1997; Shankar *et al*. 2020b; Wolf *et al*. 2020), within species (Schleucher 2004; Shankar *et al*. 2019, 2020b; Spence & Tingley 2021), and even within individuals (e.g. Krüger *et al*., 1982; Lasiewski, 1963). Broadly speaking, meta-analytic perspectives suggest that body mass explains some interspecific variation in how torpor is used across species (Geiser 1998; Ruf & Geiser 2015; Spence & Tingley 2021), as does shared ancestry: in hummingbirds, some characteristics of torpor are similar among close relatives and show phylogenetic signal (Wolf *et al*. 2020). Nevertheless, the results of evolutionary studies of natural torpor in hummingbirds have yet to be reconciled with the extensive intraspecific, ecologically driven variation in torpor, as it is challenging to meaningfully capture variation in experimental conditions across studies in meta-analyses. Hence, systematically collected data on traits of multiple species are needed to disentangle ecological from evolutionary correlates of torpor.

In the Andes, the composition of avian communities varies dramatically across elevational gradients (Quintero & Jetz 2018; Montaño-Centellas *et al*. 2020). In hummingbirds, community composition is highly phylogenetically structured (Graham *et al*. 2009), e.g. the hermit clade (Phaethornithinae) is largely absent from high elevations (Stiles 2004), which suggests that the capacity to colonize and adapt to high elevations is unequally distributed across the hummingbird phylogeny. Whether interspecific variation in the ability to use torpor contributes to interspecific variation in elevational distribution is unclear. Despite its potential importance for birds’ survival in challenging thermal environments, daily torpor is known only from 43 hummingbird species of mostly North American or extreme highland species (summarized in Spence and Tingley 2021); major data gaps exist for tropical lowland and mid-elevation species. Additionally, natural torpor has largely been measured in populations at single field sites, which prevents the sampling of environmental drivers of torpor across species’ elevational ranges.

We present the largest dataset on natural torpor in hummingbirds, both with respect to sampling effort (249 individuals) and taxonomic coverage (29 species), from a 1,920 m elevational gradient in the Western Andes of Colombia. Our data provide a unique opportunity to disentangle the relative contributions of ecological and evolutionary variation in aspects of torpor. After controlling for ecological factors, the remaining phylogenetically determined propensity to use torpor was significantly correlated with macroecological patterns of elevational range in the Andes. Evolutionary models across all Andean hummingbirds revealed numerous shifts into novel elevational ranges. Taken together, our results suggest that increases in the propensity of hummingbirds to use torpor may have played a key role in the colonization history of harsh, high-elevation Andean habitats.

## Methods

More information can be found in the supplementary methods. Briefly, we used mist nests to capture 249 individuals of 29 species of hummingbirds. In August-October 2016 and April-May 2017, we caught birds at three field sites in the western Andes of Valle del Cauca, Colombia. These sites spanned a 1,920m elevational gradient (median T_a_: 8.7 °C at 2,500 m asl to 28.4 °C at 280 m asl; Fig. S1). We caught birds in the afternoon (14:00-17:00), fed them hourly with artificial nectar (25% sucrose) through a plastic 100 mL syringe until satiation, and transferred each to an individual testing chamber (Fig. S2) at 18:00, where they remained for the entire night (∼12h). In this testing chamber, we measured T_b_ every ten seconds using cloacal thermistors, which were inserted 4-6mm and secured to the central rectrices with liquid silicone (Ricklefs & Williams 2003). T_a_ was measured simultaneously at the bottom of the chamber with a separate thermistor. Each bird was tested once, weighed immediately before entry into and upon exit from the testing chamber, fed in the morning after the night of testing until satiation, marked, and released. Torpor was entered into unambiguously in all but six trials, and we identified torpid birds based on profound decreases in T_b_ that approached T_a_ (Fig. S3) as well as on mean and variance of the frequency distribution of differences between T_a_ and T_b_ (Fig. S4).

### Torpor characteristics

For birds that used torpor, we calculated torpor bout duration (TBD) in three complementary ways that explore tradeoffs between data quality and quantity (Fig. S5). To be conservative, we present all results in the main text but focus on the unambiguous and well-sampled short TBD. We also calculated the rate of exit from torpor (Fig. S6) and the time torpor was initiated relative to local sunrise using the R package suncalc (Thieurmel & Elmarhraoui 2019). Finally, we defined the nightly loss of body mass as the difference between initial (M_b_) and final body masses (M_f_). Because the 29 species we sampled span a 5.3-fold difference in M_b_ (range: 2.93 - 15.54 g), we divided the loss of body mass by M_b_ to obtain the relative loss of body mass (%).

### Body condition

Given its constraints, tarsus length may not be an appropriate indicator of body size for Trochilidae (Fig. S7); accordingly, we consulted a separate database of 1,035 hummingbirds of the species in our study collected by us over six years of mist-netting efforts in Colombia. We discovered that within adults of well-sampled species, morphological measurements are rarely significantly correlated with body mass (tarsus: 0/17 species, bill length: 1/21 species, wing length: 10/21 species), whereas correlations were significant only across species (Fig. S8). As we had not recorded morphological measurements from the birds in our trials, we exploited these weak intraspecific allometries between morphological body size and body mass and calibrated an appropriate proxy for body condition (difference between M_b_ and species-average body mass; hereafter referred to as ‘body condition’) for the species in our sample (supplementary methods, Figs. S9-13).

### Sustained temperature gradient

Recent studies of torpor have revealed that torpor can be either “deep” or “shallow” (Mckechnie & Lovegrove 2002; Schleucher 2004; Ruf & Geiser 2015; Shankar *et al*. 2022). We captured this two-dimensional variation by approximating the area of the integral of the function of T_b_(t)-T_a_(t) between timepoints t_1_ (onset of recording) and t_2_ (end of recording) and dividing the integrated area by the temporal duration from t_1_ to t_2_ (Fig. S14). This quantity (hereafter ‘sustained temperature gradient,’ STG) describes, for a given time period, to what extent the focal bird sustained a relatively high or low difference between its internal body temperature and the environmental temperature.

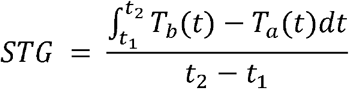

By varying t_1_ and t_2_, we calculated the STG during the pre-torpor phase (Fig. S14A) and the combined pre-torpor and torpor phases (Fig. S14B). Because the STG over the entire night (Fig. S14C) was significantly positively correlated with % loss of body mass (t = 4.297, df = 180, P < 0.0001, ρ = 0.305), we assume that STG reflects at least some portion of birds’ nightly energy expenditure (Bech *et al*. 1997; Wolf *et al*. 2020). However, we readily acknowledge that factors such as within-trial variation in thermal conductance may mask some of the signal of energy expenditure in T_b_ (M. Chappell and F. Geiser, personal communication), and that fat metabolism may also contribute to energy expenditure during torpor (Powers 1991), even if it is not a main driver of body-mass loss (c.f. Wolf *et al*., 2020). To be certain that measurement errors due to intermittent cable failure did not contaminate our estimates of STG, we calculated it only in a subset of trials with qualitatively “clean” data (N = 192). Finally, we consider the limitations of the use of STGs in the context of interspecific variation in thermal conductance and T_lc_ (Figs S15; supplementary methods).

### Statistical analysis

We investigated the drivers of torpor, and the variation in its characteristics and consequences, using phylogenetic hypotheses (Fig. S16) and Bayesian phylogenetic generalized linear mixed models (PGLMMs) implemented in the brms package (Bürkner 2017, 2018). We first examined patterns of birds entering torpor by fitting logistic PGLMMs with body mass, T_a_, body condition, and pre-torpor STG (Fig. S14A) as predictor variables, binary torpor usage as the response variable, and phylogenetic random intercepts. We also considered the effect of adding the raw data from a recent report of torpor in Andean hummingbirds at high elevations (3,800m asl; Wolf *et al*. 2020), for which we calculated the pre-torpor STG as earlier.

To evaluate drivers of variation in characteristics of torpor, we focused only on the trials that used torpor and fitted models with TBD as the response variable, using body mass, T_a_, body condition, and the STG during pre-torpor and torpor phases (Fig. S14B) as predictor variables. As practitioners define TBD in different ways (see above), we repeated this analysis for all three definitions of TBD (short, complete, and incomplete). To be conservative, we focused on short TBD as this definition strikes a balance between data quantity and quality, and we emphasize only those results that were robust to TBD definition and significant across short TBD and at least one other definition of TBD. We then calculated and analyzed variation in the rate of exit from torpor, time when birds began leaving torpor (with respect to local sunrise), and minimum T_b_ during torpor as separate response variables using the same modelling approach.

Last we analyzed the impact of torpor in two complementary analyses that used body-mass loss as the response variable and included torpor use, body condition, body mass, T_a_, and STG (across the entire night; Fig. S14C) as predictor variables. In the first analysis, we used all trials and added torpor usage as a binary predictor variable. In the second analysis, we focused on the birds that used torpor, employed short TBD as a predictor variable instead of torpor use, and considered the slope of the exit from torpor as a predictor.

For the analysis of torpor usage, the phylogenetic random intercepts represent the species-specific deviation from the average torpor usage, while holding constant the variation that is due to the remaining ecological predictor variables (these random effect levels are hereafter referred to as ‘phylogenetic torpor propensity’). We extracted these species-specific random effect levels and determined their phylogenetic signal by calculating Pagel’s λ and Blomberg’s K in phytools (Revell 2012). Finally, we synthesized elevational range data in the Andes from Quintero & Jetz (2018) and used species-level, phylogenetic regressions (PGLS) to test for a relationship between the propensity for species to enter torpor and the midpoint of their elevational range, while addressing phylogenetic uncertainty in sensiPhy (Paterno *et al*. 2018).

We then employed a process-based evolutionary model to investigate how changes in propensity for torpor influence the rate at which hummingbirds’ elevational ranges evolve (Hansen *et al*. 2021). As torpor propensity was available only from the 29 species in our study, we expanded the taxonomic scope of our data and fitted heterogeneous models of elevational range evolution with reversible jump Markov Chain Monte-Carlo (rjMCMC) to address the colonization of Andean highlands across 206 Trochilidae, using the bayou package (Uyeda & Harmon 2014). We fitted a multi-optimum rjMCMC model of elevational midpoint in bayou, using empirical priors on elevation and recommended priors on shift numbers (Uyeda *et al*. 2017). We verified model convergence (see supplementary methods) and localized well-supported shifts (posterior probability > 0.55) to different clades.

## Results

Torpor was widespread across species (used by 26/29 species) but variable across all individuals (used by 109/209 individuals in our analytic dataset). Reliance on torpor also varied substantially within and across clades (Fig. S17): 47.1% ± 0.14 (range = 6-100%, 61 individuals, 6 species) of species in the lowland hermit clade (Phaethornithinae) used torpor, while 77.8% (range = 31-100%, 80 individuals, 9 species) of the highland clade Lesbiinae used torpor. Torpid hummingbirds sustained lower temperature gradients than normothermic birds over the whole night (Fig. S14C), and torpid hummingbirds revealed short and long as well as deep and shallow torpor (Fig. S14B, S17-18).

### Torpor usage

Large birds entered torpor more often but only in response to cold ambient temperatures (Fig. 1, Table S1, Fig. S19). The effect of initial body mass on torpor usage was strong and significantly negative at cold, high-elevation sites (< 20 ºC, Farallones: posterior median: -2.047; 95% CI: -3.753 to -0.748; Fig. 2B) and non-significant at the low-elevation site (> 20 ºC, Anchicayá: posterior median: -0.247; 95% CI: -1.130 to 0.633; Fig. 2C). Body condition had a marginal, positive effect on torpor (Table S1; Fig. 2D), and birds that sustained higher pre-torpor temperature gradients significantly decreased their use of torpor (Table S1; Fig. 2E). Adding the trials from Wolf *et al*. (2020) to our analytic dataset prohibited evaluating the effect of body condition and instead confirmed strong negative effects of T_a_ and pre-torpor STG (eliminating the interaction between temperature and body mass); these trials point to a non-significant negative effect of body mass on torpor (Fig. S20).

**Figure 1.**
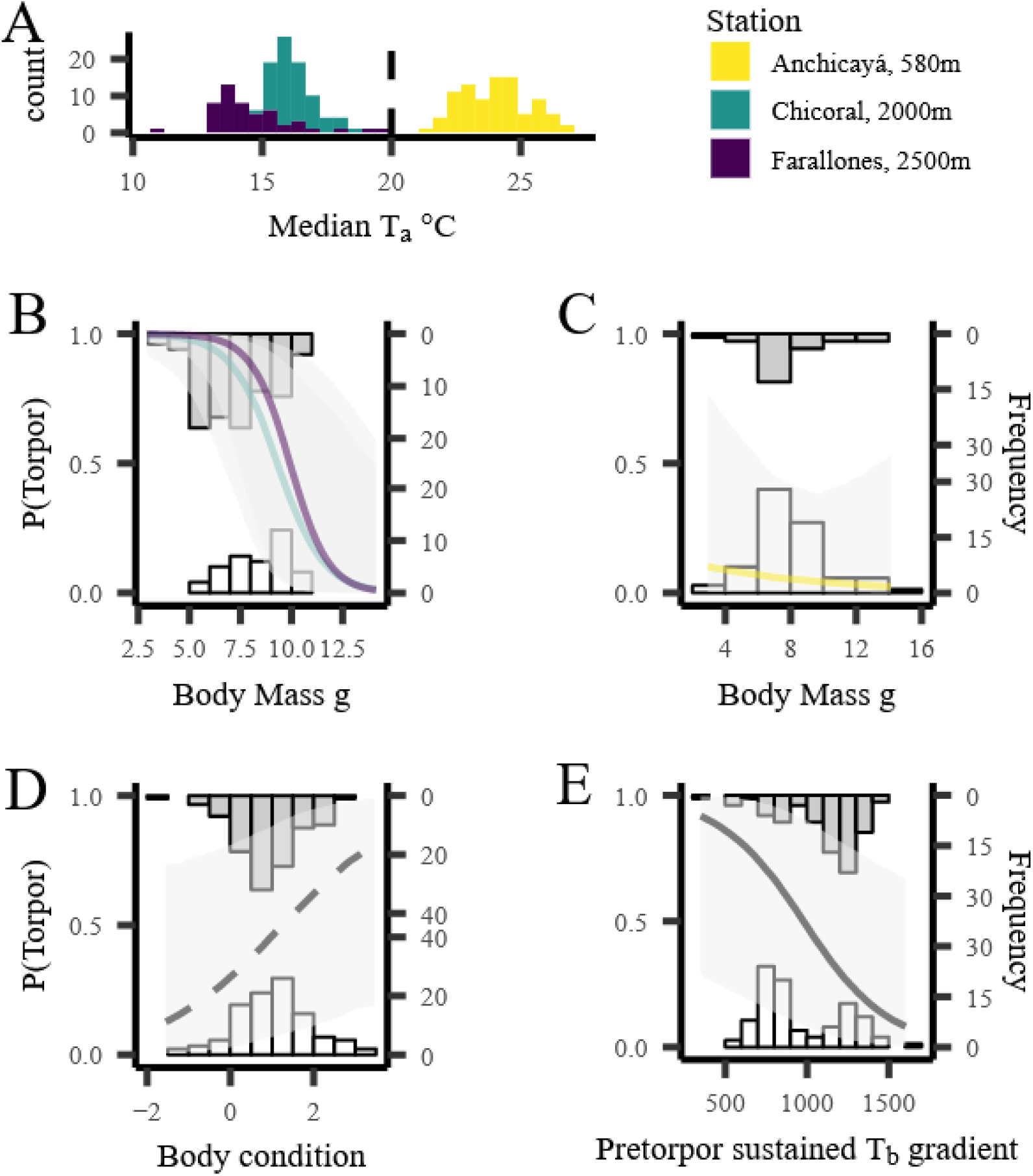
High body mass reduces the use of torpor only in cold temperatures. **A**) Frequency distributions show median T_a_ (N = 210) for each trials separated by stations. Vertical dashed line indicates 20 ºC. In **B**-**E**, histograms show frequency distributions of body mass of torpid (gray, upper) and euthermic (white, lower) birds. **B**) Significant influence of body mass on torpor below 20 ºC (at Farallones and Chicoral). **C**) Above 20 ºC (at Anchicayá), body mass had no significant effect on torpor. **D**) Body condition had marginally significant positive effect on torpor. **E**) Pre-torpor STG had a significant negative effect on torpor. Lines show conditional effects and shading shows 95% credible interval (dashed line: marginally significant; solid line: significant).

**Figure 2.**
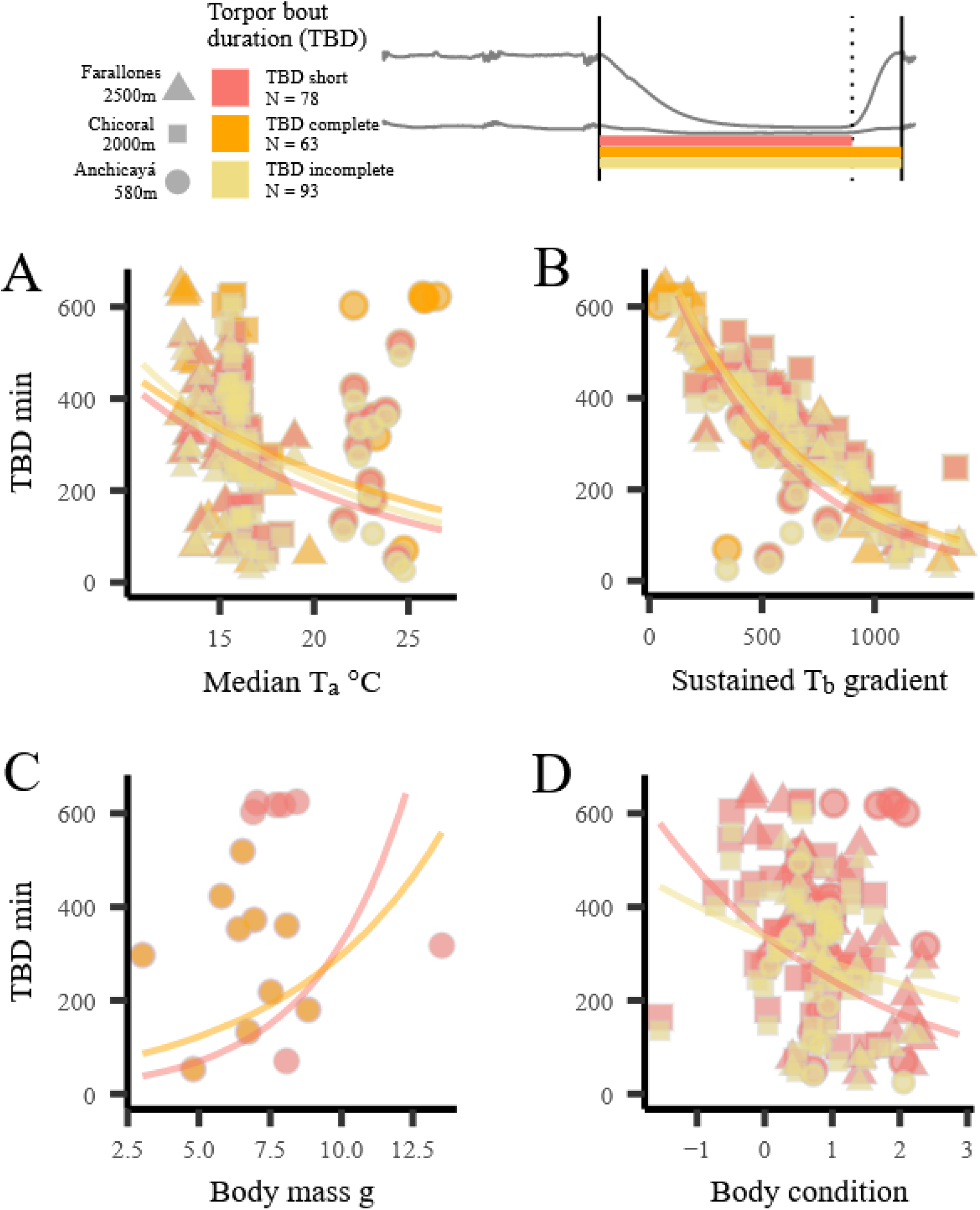
Prolonged torpor bouts reduce STG and occur when temperatures are cold and body condition is poor. Top panel shows TBD definitions on typical T_a_ and T_b,_ and vertical lines show beginning of exit from torpor (dashed), beginning of entry into and end of exit from (solid) torpor. TBD decreases significantly with increasing T_a_ (**A**) and STG (**B**) across all TBD definitions. In short and complete TBDs, a positive interaction between body mass and T_a_ yielded a positive effect of body mass on TBD at T_a_ > 20ºC (**C**). Body condition had a significant negative effect on short and incomplete TBDs (**D**).

### Torpor characteristics

Across all TBD definitions, torpor tended to be short and shallow when T_a_ was warm, yielding high STG values. When T_a_ was cold, torpor was prolonged and deep, yielding low STG values (Table S2, Fig. 2A-B, S21-23). Moreover, in both short and complete TBD, there was a consistent, significant, positive interaction between T_a_ and body mass (Table S2, Fig. S21-23): at cold, high elevations, TBD was unaffected by body mass (Farallones: short TBD posterior median of body mass: -0.102, 95% CI: -0.296 – 0.070), but at the warm, low-elevation station, birds with higher body mass remained longer in torpor (Anchicayá; short TBD posterior median of body mass: 0.325, 95% CI: 0.048 – 0.612; Fig. 2C, S21). The better a bird’s body condition, the less time it spent in torpor, according to definitions of both short and incomplete TBD (Table S2; Fig. 2D, S21-23).

Torpor was exited more rapidly when T_a_ was cold and birds were small (Table S3; Fig. 3A-B, S24). However, we found a significant negative interaction between body mass and STGs, suggesting that birds that had sustained large temperature gradients returned to normothermy more slowly and irrespective of body mass (90^th^ quantile STG: posterior median of body mass: 0.055, 95% CI: -0.232 – 0.343) compared to birds that had sustained low temperature gradients (10^th^ quantile STG: posterior median of body mass: -0.344, 95% CI: -0.577 – -0.113; Fig. 3B). In cold environments, birds waited until shortly before sunrise before initiating their exit from torpor (Table S3; Fig. 3C, S25). Additionally, small birds began leaving torpor earlier at night than large birds, but only when body condition was high (90^th^ quantile body condition posterior median of body mass: -0.205, 90% CI: - 0.402 – -0.013; Table S3, Fig. 3D). Birds with low body condition exited torpor irrespective of body mass (10^th^ quantile body condition posterior median of body mass: - 0.010, 95% CI: -0.174 – 0.151).

**Figure 3.**
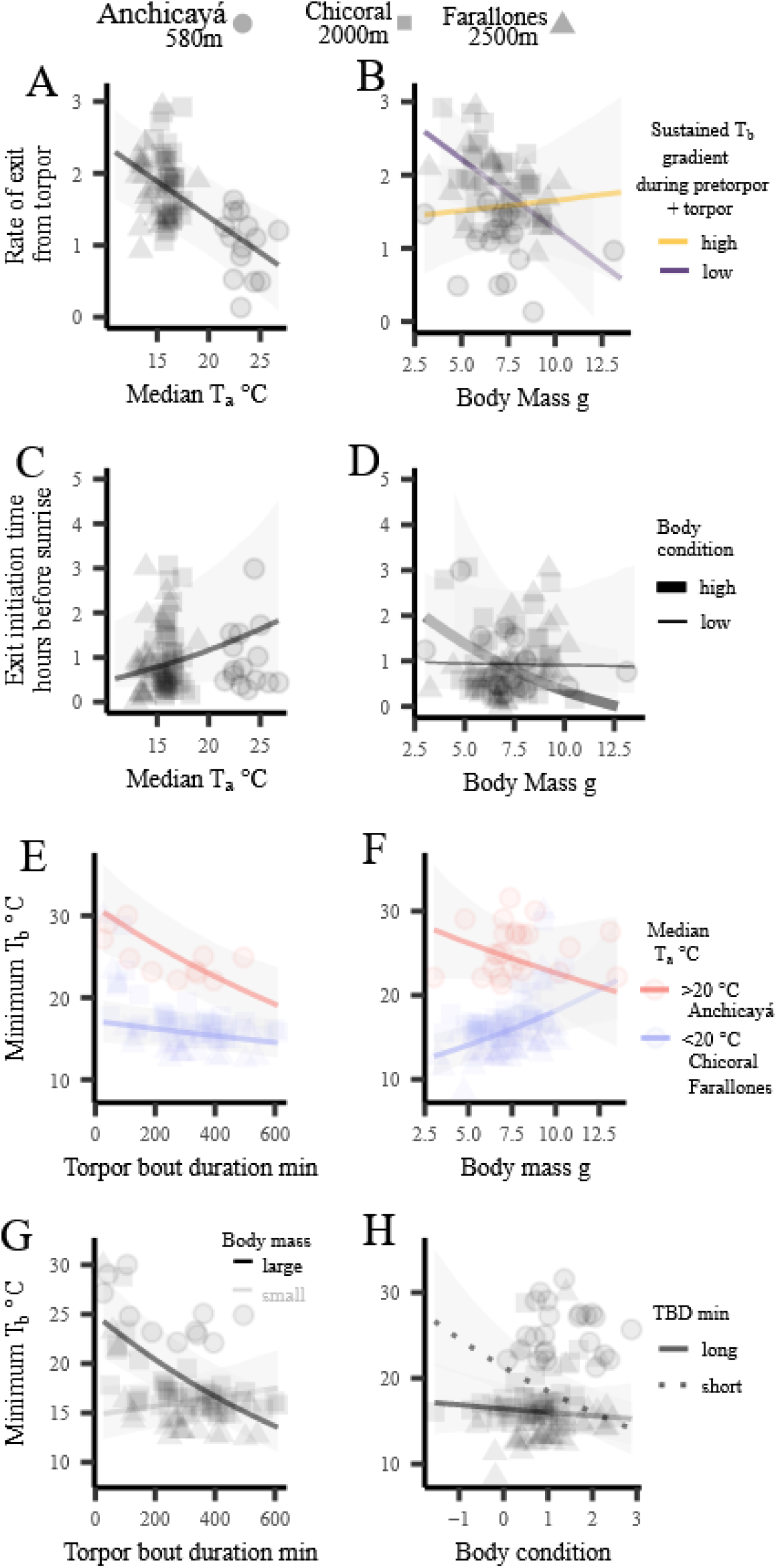
Internal and external factors govern the rate and time of exit from torpor as well as torpor depth. Rate of exit from torpor decreases as T_a_ (**A**) and body mass (**B**) increase. Time of exit from torpor increases with T_a_ (**C**) and decreases with body mass (**D**). Interactions in **B, D, G** and **H** are plotted with 10^th^ and 90^th^ quantiles. Lines show conditional effects and shading shows 95% credible intervals.

During torpor, minimum T_b_ was chiefly influenced by T_a_ (Table S3; Fig. 3E-F, S18, S26) but was also significantly influenced by body condition, body mass, and STG, as well as by other complex interactions (Table S3; Fig. 3E-F). Large birds (90^th^ quantile body mass posterior median of TBD: -0.146, 95% CI: -0.199 – -0.091) decreased minimum T_b_ by prolonging TBD (10^th^ quantile body mass posterior median of TBD: 0.041, 95% CI: - 0.015 – 0.100; Fig. 3G). Additionally, the minimum T_b_ of birds was unaffected by body condition during long torpor bouts (90^th^ quantile TBD posterior median of body condition: - 0.019, 95% CI: -0.082 – 0.045). But for birds that relied on short torpor bouts, the better a bird’s body condition, the deeper its torpor (10^th^ quantile TBD posterior median of body condition: -0.010, 95% CI: -0.178 – -0.050; Fig. 3H).

### Torpor impact

Across all birds, relative body-mass loss was substantial (17.1% ± 0.01, range = 0-32.4%, N = 237 individuals, N = 28 species). The larger the bird, the less body mass it lost (posterior median of body mass: - 0.020; 95% CI: -0.037 – -0.006; Table S4; Fig. 4A, S27). Although torpor usage itself did not reduce loss of body mass (posterior median: -0.006, 95% CI: -0.026 – 0.012), it decreased the STG (Fig. S14C), which in turn avoided losing mass (posterior median for effect of STG: 0.013; 95% CI: 0.001 – 0.022; Fig. 4B). Birds with better body condition lost more relative body mass (Fig. 4C). However, in birds that used torpor, body condition affected the loss of body mass more (by a factor of 1.9; posterior median of body condition: 0.054; 95% CI: 0.038 – 0.070) than in birds that did not use torpor (posterior median of body condition: 0.029; 95% CI: 0.014 – 0.043).

**Figure 4.**
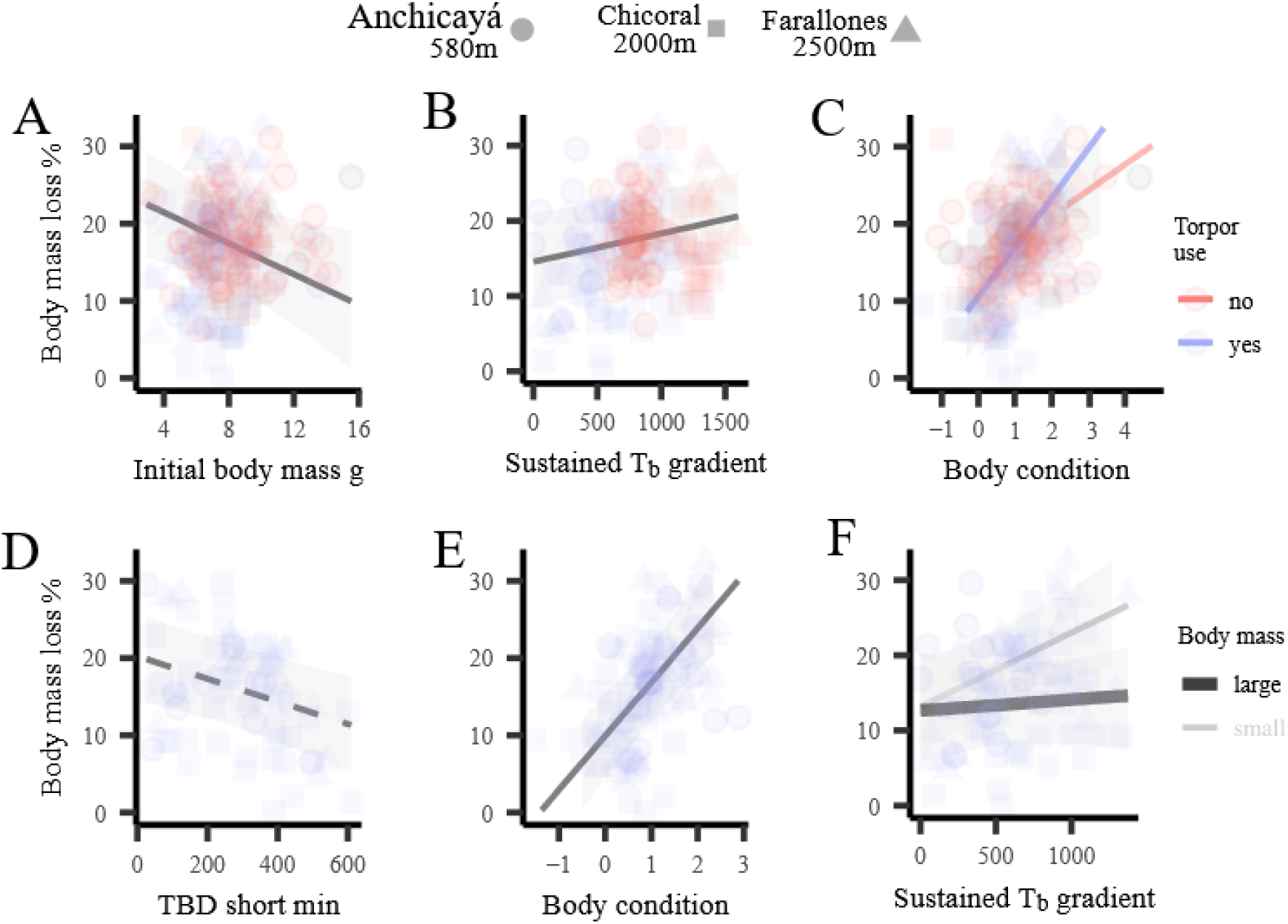
Internal and external factors govern the impacts of torpor. Relative mass loss across all birds (**A-C**) declines with initial body mass (**A**), increases with increasing STG (**B**) and increases with body condition, albeit more steeply for torpid birds. Within torpid birds (**D-F**), relative body-mass loss declines with longer torpor when STG is not considered (**D**), increases with body condition, (**E**) and increases with higher STG, although only for birds with small body mass (**F**). Interaction and **F** is plotted with 10^th^ and 90^th^ quartiles. Lines show conditional effects and shading shows 95% credible intervals.

In birds that used torpor, we found a significant negative effect of TBD on relative body-mass loss in the full dataset (posterior median for effect to TBD: -0.021; 95% CI: - 0.036 – -0.009; Fig. 4D, S28). However, once we considered STG as a predictor variable in the reduced dataset, this effect was rendered not significant (Table S4, Fig. S28), likely because longer TBD contributed to reduced STG. Body condition had a positive effect on relative body-mass loss (posterior median for effect to body condition: 0.053; 95% CI: 0.036 – 0.068; Fig. 4E). The positive effect of the STG on body-mass loss was modulated by initial body mass (Table S4, Fig. 4F). Large birds lost body mass irrespective of how large of a temperature gradient they sustained (body mass 90^th^ quantile, posterior median of STG: 0.004; 95% CI: -0.018 to 0.025, Fig. 4F). In contrast, small birds lost 7.4 times more body mass per unit of temperature gradient sustained (body mass 10^th^ quantile, posterior median of STG: 0.030; 95% CI: 0.012 to 0.049, Fig. 4F).

### Evolutionary consequences of torpor for elevational changes

Of all traits associated with torpor we analyzed, phylogeny contributed most strongly to the probability of deploying torpor; a considerable portion of its variance was attributable to phylogenetic random effects (posterior median = 0.29; Table S5) and phylogenetic signal for torpor usage was high (posterior median λ = 0.96; Table S6). For all other characteristics of torpor, between 10 and 33% of the variation in time spent in torpor was attributable to intraspecific variation, and much less (<2%) variation was due to phylogeny (Table S5). Our model of torpor usage controls for ecological sources of variation and the phylogenetic random intercepts describe the species-level propensity for torpor. This propensity significantly explained variation in elevational midpoints across the Andes in standard (linear model, torpor propensity: 622.3 ± 238.6, T = 2.61, P = 0.015, Fig. 5A dashed line) and phylogenetic regression models (PGLS, torpor propensity: 699.2 ± 313.32, T = 2.23, P = 0.034, Fig. 5A, solid line), as well as regression with phylogenetic uncertainty (torpor propensity coefficient: 692.2 ± 310.4, P = 0.034). However, our limited taxonomic sampling suggested that the correlation was chiefly driven by species from the highland clades Lesbiini (coquettes) and Heliantheini (brilliants, Fig. 5).

**Figure 5.**
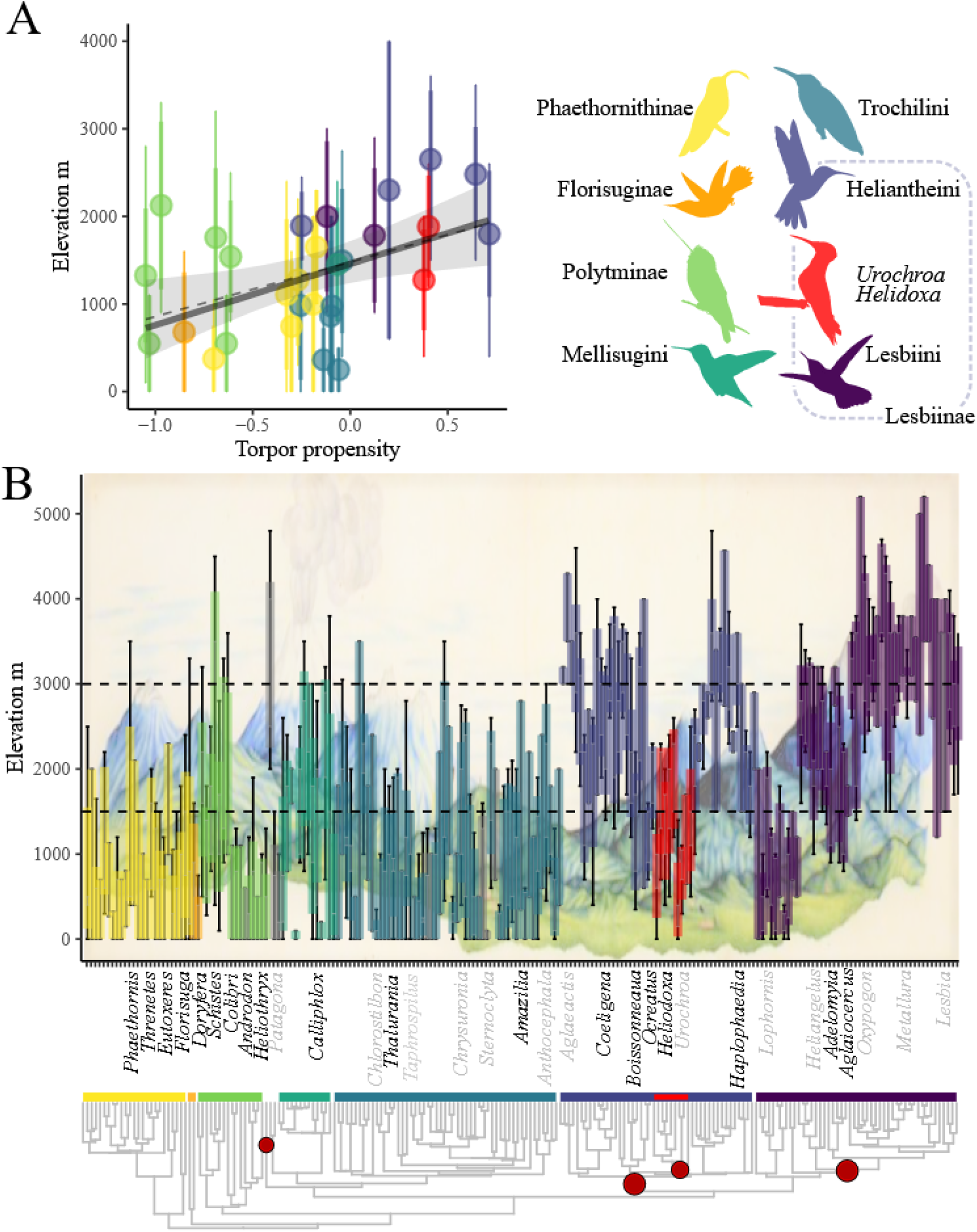
Variation in the propensity for torpor matches elevational range distributions across hummingbird clades. **A**) Phylogenetic propensity for torpor significantly increases elevational midpoints in both linear (dashed line) and phylogenetic regression models (solid line). Points show elevational midpoint across the Andes. Vertical bars show mean (thick) and range (thin) of elevational limits. Gray shading shows 95% confidence intervals from PGLS regression. **B**) Shows results of rjMCMC model of evolution of elevational midpoints across 206 Andean hummingbirds. Significant, robust (posterior probability > 0.55) shifts in elevation are denoted by red circles, whose diameter is proportional to posterior probability. Genera shown in black are represented in our study of torpor. Panel background in B shows altitudinal stratification in the Andes from Caldas (1803).

The process-based models of how the propensity for torpor affected the rate of elevational evolution revealed low absolute phylogenetic R^2^ (6.03%), but with a notable mean effect: the predicted rate variance, in units of squared (scaled, centered, and positive) meters in elevation per tree length (h = 27.15 my), is 0.93 + 0.77 x, where x (scaled, centered, and positive) is the propensity for torpor. This means that the 1.85-fold difference between the propensity of hermits to use torpor (Phaethornithinae mean = -0.33) and brilliant/coquette torpor propensity (Lesbiinae mean = 0.28) would elicit a 68.7% increase in variance in the rate of evolutionary change in elevation, i.e., a (√ 1.687) 29.9% increase in elevational change rate. However, unsurprisingly (given the low sample size, N = 29), this effect size revealed high uncertainty and was not significant (bootstrapped SE = 0.58; 95% confidence interval: -0.639 – 2.259). We then modeled evolutionary changes in elevation across 206 Andean hummingbirds with rjMCMC multi-optimum models. The model converged satisfactorily (all □ < 1.104; Table S7, Fig. S29) and revealed strong support (posterior probabilities > 55%) for four shifts in elevational optima (Table S8). A null model with constrained numbers of shifts in elevation failed to recover these shifts (Table S9). Colonization of the Andean highlands appeared in the highland specialist Giant Hummingbird (*Patagona gigas*) and in 14 genera within the highland Lesbiinae (Lesbiini and Heliantheini; Table S8; Fig. 5B). Strikingly, we also discovered a well-supported descent in elevation into low/mid elevations by two genera (*Heliodoxa* and *Urochroa*) that are nested within the highland Heliantheini (Table S8; Fig. 5B).

## Discussion

Our current dataset almost doubles the current taxonomic coverage of torpor studies in hummingbirds, and reveals wide variation in the use of torpor, its characteristics and impacts, both within and across species, along a 1,920 m tropical elevational gradient. We find that external conditions, chiefly T_a_, drive the characteristics of torpor: cold temperatures made birds more likely to deploy (Fig. 1A-C) and prolong torpor (Fig. 2A), and caused birds to accelerate their departure from torpor (Fig. 3A), delay their departure from torpor until immediately before sunrise (Fig. 3C), and rapidly thermoconform. The duration of an individual’s torpor (Fig. 3E) and its body mass (Fig. 3F) were governed by complex effects. These findings add to a growing body of literature that suggests hummingbirds use numerous external cues to decide if, when, and how to deploy torpor (e.g. Calder and Booser 1973; Eberts *et al*. 2021; Shankar *et al*. 2019).

Our results also reveal high intraspecific variability in torpor’s characteristics, with important roles played by proxy variables for internal individual-level conditions: energy reserves (body condition) and energy expenditure (STG). Birds with high body condition were marginally more likely to use torpor (Fig. 1D) than birds with low body condition, and when they did, displayed short TBDs (Fig. 2D), which allowed them to regain normothermy well before sunrise; this effect was particularly important for small-bodied species (Fig. 3D). Birds with high body condition lost more mass over the course of the night compared to birds with low body condition (Fig. 4C). As the effect was strongest for torpid birds (Fig. 4C), we suspect that body condition captures at least some of the energy stores that birds use to fuel the costly return to normothermy from a torpid state. Although many studies have emphasized the energetic benefits of torpor (Calder & Booser 1973; Carpenter 1974), we suggest that torpor has potentially substantial costs as well, perhaps chiefly in the form of the return to normothermy and the maintenance of normothermy before sunrise (Shankar *et al*. 2020b; Eberts *et al*. 2021). These costs place particular premiums on passive warming strategies such as solar “hitch-hiking” (Wolf *et al*. 2020) and may factor into individual birds’ decisions to enter into torpor. Entering into torpor without the necessary energy to depart may mean birds forgo important early-morning foraging opportunities when nectar crops are high and not yet depleted by competitors (Paton & Carpenter 1984); alternatively, they may find exiting torpor insurmountable without assistance (Wolf *et al*. 2020). Overall, when faced with cold, nocturnal T_a_, hummingbirds must choose between two risky options: gambling on maintaining normothermy, or deploying and attempting to recover from torpor.

Calder and Booser (1973) described hypothermic hummingbirds as possessing a “biological fuel gauge” that indicates energy stores and consumption. Although we measured only a coarse proxy of energy consumption (STG, Fig. S14), we nevertheless found STG influences different aspects of torpor. Birds that sustained high temperature gradients early in the night were less likely to enter torpor than those that sustained low-temperature gradients. We suspect that this correspondence may mean hummingbirds enter a shallow phase of torpor before committing to a full drop in metabolism (Shankar *et al*. 2022), which decreases pre-torpor STG. Additionally, prolonging torpor decreased STG during the pre-torpor and torpor phases (Fig. 2B, S14B), yielding an obvious energetic benefit (Shankar *et al*. 2020b). Interestingly, birds with high STG exited torpor slowly, regardless of body mass (Fig. 3B), whereas small birds that had incurred low STG values showed the fastest rates of exit from torpor. This interaction suggests that birds that had failed to minimize energy expenditure over the night are forced to rewarm more slowly, which might foreclose important early-morning foraging opportunities (Paton & Carpenter 1984). Moreover, we showed that not only torpor duration (Shankar *et al*. 2020b), but the combined depth and duration (as captured by STG over the entire night, Fig. S14C) was crucial to predicting variation in nightly body-mass loss (Fig. 4B). Finally, the effect of STG varied predictably across small and large species: torpid birds that sustained large temperature gradients incurred large nightly body-mass loss, but small birds lost more than large birds per unit of STG exposure (Fig. 4F). This finding matches fundamental theoretical predictions about the role of body size in thermoregulation, as large individuals should suffer less from being exposed to punishing thermal gradients (and pay fewer costs in terms of body-mass loss) than small individuals, due to decreased ratios of surface area to volume (Bergmann 1847).

We confirmed previous claims that natural torpor usage is influenced by body mass (Spence & Tingley 2021), but we found that this effect holds true only for torpor in cold environments. Sampling within species across elevational gradients was necessary to reveal this context-dependent effect of body mass on natural torpor, as adding data from trials conducted at a single, extremely cold high-elevation site (3,800m asl, 2.4 - 5.9°C; Wolf *et al*. 2020) erased the effect of body mass (Fig. 1, S19). All individuals in that study entering torpor, even the individuals of the largest hummingbird species, the Giant Hummingbird (*Patagona gigas*, mean adult body mass: 20.2 g). Thus, extreme climatic harshness may reduce variation in strategies birds use to enter torpor, suggesting that torpor is required to survive in the highest habitats in the Andes. But at low and intermediate elevations (< 2,500 m asl), we documented effects of body condition and pre-torpor STG, which emphasize the importance of individual-level energy budgets that hummingbirds may consider when deciding to enter torpor (Calder & Booser 1973).

The substantial intraspecific variability in all aspects of torpor (deployment, characteristics and impact) further supports the idea that entering torpor is a highly variable strategy that is judiciously deployed and modulated, depending on internal and external conditions. This variation contrasts with claims about evolutionary variation in torpor, most of which use single-species averages that are frequently derived from measurements at single study sites (Wolf *et al*. 2020; Spence & Tingley 2021). In our study, most variance in traits associated with torpor was intraspecific (Table S5), and the phylogenetic signal for these traits was generally low (Table S6), with one exception: propensity for torpor. Our measure of propensity for torpor accounts for its extensive underlying ecological variation, so it remains an informative species-level trait. The phylogenetic propensity for torpor varied across clades and with macroecological patterns of elevational range (Fig. 5), and the resulting discontinuous pattern matches the strong phylogenetic community structure of hummingbirds across elevations (Graham *et al*. 2009). Despite not achieving significance, the process-based model of torpor propensity’s effect on the rate of change in elevational midpoints suggests a trend where clades with a low propensity for torpor (e.g. Polytminae, Phaethornithinae) should show few elevational changes, whereas clades with a high propensity (e.g. Lesbiinae) show frequent, and even bidirectional, elevational shifts. In line with this prediction, we localized robust upslope and downslope shifts precisely in the clades that tended to have a high propensity for torpor (Lesbiinae, Fig. 5B, Table S7-8). Our results are consistent with the general idea that the propensity for torpor in hummingbirds might function as a key innovation (Simpson 1944), although more data are required for a rigorous test (Miller & Stroud 2022). When Stiles (2004) asked, “why are there no hermits in the paramo?” he was “unaware of any studies on physiology and torpor in hermits.” Our study adds to at least one other report of torpor in hermits (Shankar *et al*. 2020a), but discrete reports alone are unlikely to illuminate family-wide elevational patterns. Instead, our results suggest that continuous variation in the ecologically informed propensity for torpor may have facilitated Lesbiinae’s colonization of and proliferation within the harsh climates of the Andes.

## Acknowledgements

We are grateful to many people for their help during fieldwork: Maria Fernanda Restrepo, Kirsty Macphie, Lina Peña, Mariana Cortés, Alexandra Buitrago, Mario Agustín Loaiza, and Karen Tatiana Niño, for their invaluable support during long sleepless nights, and Betty Cadena (La Minga Ecolodge), Juan Vicente (Zygia Biological Station), and all the staff of EPSA-Anchicayá, for their affection, hospitality, and assistance at the sampling sites. We also thank Dr. Jose Miguel Chaves-Fallas and Dr. Molly Womack for help with figure design; Aryeh Miller and Sean McHugh for insights on evolutionary analyses. Dr. Jimmy McGuire kindly provided a phylogenetic tree file. Dr. Mark Chappell and Dr. Fritz Geiser shared fruitful discussions about torpor, and the following contributors to phylopic.org for sharing their images: Ferran Sayol, Mike Keesey, Margot Michaud, Emma Hughes, and Steven Traver. Finally, we are grateful to Icesi University (Zygia Biological Station), La Minga Ecolodge, PNN Farallones de Cali (granted the permits), and Celsia (Anchicayá Station) for allowing us to work at the different field sites. This research was funded by a National Science Foundation Grant (DEB-1120682), an Alexander Skutch Award (Association of Field Ornithologists), a Louis Agassiz Fuertes Award (Wilson Ornithological Society), and an Alexander Wetmore Award (American Ornithologists’ Union) to GAL and by a Fulbright Student Research scholarship to JWB.

## References

Altshuler, D.L., Dudley, R. & McGuire, J.A. (2004a). Resolution of a paradox: Hummingbird flight at high elevation does not come without a cost. Proc. Natl. Acad. Sci. U. S. A., 101, 17731–17736.

Altshuler, D.L., Stiles, F.G. & Dudley, R. (2004b). Of Hummingbirds and Helicopters: Hovering Costs, Competitive Ability, and Foraging Strategies. Am. Nat., 163, 16–25.

Bech, C., Abe, A.S., Steffensen, J.F., Berger, M. & Bicudo, J.E.P. (1997). Torpor in three species of Brazilian hummingbirds under semi-natural conditions. Condor, 780–788.

Bergmann, C. (1847). Über die Verhältnisse der Wärmeökonomie der Thiere zu ihrer Größe. Göttinger Stud., 1, 595–708.

Bürkner, P.C. (2017). brms: An R package for Bayesian multilevel models using Stan. J. Stat. Softw., 80.

Bürkner, P.C. (2018). Advanced Bayesian multilevel modeling with the R package brms. R J., 10, 395–411.

Caldas, J.F. de. (1803). La Memoria sobre la Nivelación de las Plantas del Ecuador, Historia de Nuestra Revolución, Educación de Menores, Importancia del Cultivo de la Cochinilla y Chinchografía y Geografía de los Arboles de Quina.

Calder, W.A. & Booser, J. (1973). Hypothermia of broad-tailed hummingbirds during incubation in nature with ecological correlations. Science (80-.)., 180, 751–753.

Carpenter, F.L. & Hixon, M.A. (1988). A New Function for TorporL: Fat Conservation in a Wild Migrant Hummingbird. The Condor1, 90, 373–378.

Carpenter, L.F. (1974). Torpor in an andean hummingbird: its ecological significance. Science (80-.)., 183, 545–547.

Eberts, E.R., Guglielmo, C.G. & Welch, K.C. (2021). Reversal of the adipostat control of torpor during migration in hummingbirds. Elife, 10, 1–21.

Fristoe, T.S., Burger, J.R., Balk, M.A., Khaliq, I., Hof, C. & Brown, J.H. (2015). Metabolic heat production and thermal conductance are mass-independent adaptations to thermal environment in birds and mammals. Proc. Natl. Acad. Sci. U. S. A., 112, 15934–15939.

Geiser, F. (1998). Evolution of daily torpor and hibernation in birds and mammals: Importance of body size. Clin. Exp. Pharmacol. Physiol., 25, 736–740.

Geiser, F. (2004). Metabolic rate and body temperature reduction during hibernation and daily torpor. Annu. Rev. Physiol., 66, 239–274.

Geiser, F., Currie, S.E., O’Shea, K.A. & Hiebert, S.M. (2014). Torpor and hypothermia: Reversed hysteresis of metabolic rate and body temperature. Am. J. Physiol. - Regul. Integr. Comp. Physiol., 307, R1324–R1329.

Graham, C.H., Parra, J.L., Rahbek, C. & McGuire, J.A. (2009). Phylogenetic structure in tropical hummingbird communities. Proc. Natl. Acad. Sci. U. S. A., 106, 19673–19678.

Hainsworth, F.R., Collins, B.G. & Wolf, L.L. (1977). The Function of Torpor in Hummingbirds. Physiol. Zool., 3, 215–222.

Hansen, T.F., Bolstad, G.H. & Tsuboi, M. (2021). Analyzing Disparity and Rates of Morphological Evolution with Model-Based Phylogenetic Comparative Methods. Syst. Biol., 0, 1–19.

Hiebert, S.M. (1991). Seasonal Differences in the Response of Rufous Hummingbirds to Food Restriction. Condor, 93, 526–537.

Janzen, D.H. (1967). Why Mountain Passes are Higher in the Tropics. Am. Nat., 101, 233–249.

Krüger, K., Prinzinger, R. & Schuchmann, K.L. (1982). Torpor and metabolism in hummingbirds. Comp. Biochem. Physiol. -- Part A Physiol., 73, 679–689.

Lasiewski, R.C. (1963). Oxygen Consumption of Torpid, Resting, Active, and Flying Hummingbirds. Physiol. Zool., 36, 122–140.

McGuire, J.A., Witt, C.C., Remsen, J. V., Corl, A., Rabosky, D.L., Altshuler, D.L., et al. (2014). Molecular phylogenetics and the diversification of hummingbirds. Curr. Biol., 24, 910–916.

Mckechnie, A.E. & Lovegrove, B.G. (2002). Avian Facultative Hypothermic Responses: A Review. The Condor2, 104, 705–724.

Miller, A.H. & Stroud, J.T. (2022). Novel Tests of the Key Innovation Hypothesis: Adhesive Toepads in Arboreal Lizards. Syst. Biol., 71, 139–152.

Montaño-Centellas, F.A., McCain, C. & Loiselle, B.A. (2020). Using functional and phylogenetic diversity to infer avian community assembly along elevational gradients. Glob. Ecol. Biogeogr., 29, 232–245.

Myers, N., Mittermeler, R.A., Mittermeler, C.G., Da Fonseca, G.A.B. & Kent, J. (2000). Biodiversity hotspots for conservation priorities. Nature, 403, 853–858.

Paterno, G.B., Penone, C. & Werner, G.D.A. (2018). sensiPhy: An r-package for sensitivity analysis in phylogenetic comparative methods. Methods Ecol. Evol., 9, 1461–1467.

Paton, D.C. & Carpenter, F.L. (1984). Peripheral foraging by territorial rufous hummingbirds: defense by exploitation. Ecology, 65, 1808–1819.

Pearson, O.P. (1950). The Metabolism of Hummingbirds. Condor, 52, 145–152.

Pearson, O.P. (1953). Use of Caves by Hummingbirds and Other Species at High Altitudes in Peru. Condor, 55, 17–20.

Porter, W.P. & Kearney, M. (2009). Size, shape, and the thermal niche of endotherms. Proc. Natl. Acad. Sci. U. S. A., 106, 19666–19672.

Powers, D. (1991). Diurnal variation in mass, metabolic rate, and respiratory quotient in Anna’s and Costa’s hummingbirds. Physiol. Zool., 64, 850–870.

Powers, D.R., Brown, A.R. & Van Hook, J.A. (2003). Influence of normal daytime fat deposition on laboratory measurements of torpor use in territorial versus nonterritorial hummingbirds. Physiol. Biochem. Zool., 76, 389–397.

Projecto-Garcia, J., Natarajan, C., Moriyama, H., Weber, R.E., Fago, A., Cheviron, Z.A., et al. (2013). Repeated elevational transitions in hemoglobin function during the evolution of Andean hummingbirds. Proc. Natl. Acad. Sci. U. S. A., 110, 20669–20674.

Quintero, I. & Jetz, W. (2018). Global elevational diversity and diversification of birds. Nature, 555, 246–250.

Revell, L.J. (2012). phytools: An R package for phylogenetic comparative biology (and other things). Methods Ecol. Evol., 3, 217–223.

Ricklefs, R.E. & Williams, J.B. (2003). Metabolic responses of shorebird chicks to cold stress: Hysteresis of cooling and warming phases. J. Exp. Biol., 206, 2883–2893.

Ruf, T. & Geiser, F. (2015). Daily torpor and hibernation in birds and mammals. Biol. Rev., 90, 891–926.

Schleucher, E. (2004). Torpor in birds: Taxonomy, energetics, and ecology. Physiol. Biochem. Zool., 77, 942–949.

Shankar, A., Cisneros, I.N.H., Thompson, S., Graham, C.H. & Powers, D.R. (2022). A heterothermic spectrum in hummingbirds. J. Exp. Biol., 225, 1–10.

Shankar, A., Graham, C.H., Canepa, J.R., Wethington, S.M. & Powers, D.R. (2019). Hummingbirds budget energy flexibly in response to changing resources. Funct. Ecol., 33, 1904–1916.

Shankar, A., Powers, D.R., Dávalos, L.M. & Graham, C.H. (2020a). The allometry of daily energy expenditure in hummingbirds: An energy budget approach. J. Anim. Ecol., 89, 1254–1261.

Shankar, A., Schroeder, R.J., Wethington, S.M., Graham, C.H. & Powers, D.R. (2020b). Hummingbird torpor in context: duration, more than temperature, is the key to nighttime energy savings. J. Avian Biol., 51, 1–14.

Simpson, G. (1944). Tempo and Mode in Evolution. Columbia University Press, New York, NY.

Spence, A.R. & Tingley, M.W. (2021). Body size and environment influence both intraspecific and interspecific variation in daily torpor use across hummingbirds. Funct. Ecol., 35, 870–883.

Stiles, F.G. (2004). Phylogenetic Constraints Upon Morphological and Ecological Adaptation in Hummingbirds (Trochilidae): Why Are There No Hermits in the ParamoL? Ornitol. Netropical, 15, 191–198.

Thieurmel, B. & Elmarhraoui, A. (2019). suncalc: Compute Sun Position, Sunlight Phases, Moon Position and Lunar Phase.

Uyeda, J.C. & Harmon, L.J. (2014). A novel Bayesian method for inferring and interpreting the dynamics of adaptive landscapes from phylogenetic comparative data. Syst. Biol., 63, 902–918.

Uyeda, J.C., Pennell, M.W., Miller, E.T., Maia, R. & McClain, C.R. (2017). The evolution of energetic scaling across the vertebrate tree of life. Am. Nat., 190, 185–199.

Vuilleumier, F. (1969). Field Notes on Some Birds From the Bolivian Andes. Ibis (Lond. 1859)., 111, 599–608.

Wolf, B.O., Mckechnie, A.E., Schmitt, C.J., Czenze, Z.J., Johnson, A.B., Witt, C.C., et al. (2020). Extreme and variable torpor among high-elevation Andean hummingbird species. Biol. Lett., 5–9.

Wolf, B.O. & Walsberg, G.E. (2000). The role of the plumage in heat transfer processes of birds. Am. Zool., 40, 575–584.

