## Supplementary Methods for "Ecological drivers and evolutionary consequences of torpor in Andean hummingbirds"

| Field sites……………………………………………………………...………………... | 2 |
| --- | --- |
| Bird capture and experimental protocol……………………………………………....... | 2 |
| Defining torpor……………………………………………………………..................... | 4 |
| Characteristics of torpor……………………………………………………………....... | 5 |
| Body condition……………………………………………………………..................... | 7 |
| Sustained temperature gradient…………………………………………………………. | 9 |
| Phylogenetic hypotheses……………………………………………………………….. | 12 |
| Statistical analysis – torpor experiments……………………………………………….. | 12 |
| Statistical analysis – torpor experiments with expanded dataset……………………….. | 14 |
| Statistical analysis – phylogenetic propensity for torpor and phylogenetic signal…....... | 15 |
| Elevational ranges ………………….…………………………………………………... | 15 |
| Statistical analysis – evolutionary models……………………………………………… | 16 |
| Supplementary figures …………………………………………………………………. | 19-48 |
| Supplementary tables …………………………………………………………………. | 49-60 |
| References………………………………………………………………………………. | 61-64 |

*Field sites*

The highest location where we sampled hummingbirds is the Icesi University research station, Zygia, situated on the eastern slope of the western Andes range in the Farallones de Cali National Park, Cali, (3°27’10.2” N, 76°46’28.8” W; 2,500 m). The habitat at Zygia, high-elevation Andean cloud forest, had a mean annual temperature of 14.0 °C (range = 8.7–24.8 °C) and total precipitation of 1,664 mm during 2016. The second-highest site is the La Minga Ecolodge adjacent to the Bitaco Forest Reserve, Vereda Chicoral, La Cumbre (03°33’54.2” N, 76°35’00.0” W; 2,000 m). This habitat, montane cloud forest, had an average annual temperature of 16.4 °C (range = 13.0- 22.7 °C) and total precipitation of 1315 mm during 2016. The lowest site, Anchicayá, is a tropical rainforest in the Chocó region (3°59’12.7” N, 76°88’24.4” W; 580 m), which is tropical had an average annual temperature of 21.7°C (range = 17.9 - 28.4°C) during 2016 and total annual precipitation of 5965mm. We measured temperature with dataloggers across annual cycles between 2016 and 2019 from inside the forest (Fig. S1A). We estimated slopes of T_a_ over time of day by fitting linear models to the temperature data of each station. Between sunrise near 6:00 AM and noon, T_a_ increased between 1.9 and 2.4 times faster at Farallones, the highest elevation station, than at the lower stations (Fig. S1B).

*Bird capture and experimental protocol*

Birds were captured with mist-nets during the afternoon (14:00-17:00 h). Before starting the measurements, each individual bird was fed hourly with artificial nectar (25% sucrose) until satiation. Immediately after sunset (between 18:00-19:00), each bird was weighed with a pocket scale with an accuracy of ± 0.05 g (FlipScale F2, [www.myweigh.com/pocket](http://www.myweigh.com/pocket)) and placed in a small cylindrical plastic cylinder (7cm high and 8 cm wide; ~300 ml) that contained a perch (Fig. S2). In order for the hummingbirds to experience natural field temperatures and photoperiod, the chamber lid was replaced with a mosquito net and the cylinder was placed near vegetation at night. As these birds roost nocturnally within vegetation that blocks direct radiant heat loss towards the atmosphere, we painted the outside surfaces of the chambers black to approximate this reduction in reflected radiation (Porter 1969). We monitored T_b_ and T_a_

every 10 seconds throughout the night with thermistors (2x3 mm; Measurement Specialties, Inc. part number GA10K4A80I) that were attached to a U-12 4 channel HOBO data logger (Onset Computer Corporation http//www.onsetcomp.com) that recorded the temperature. We inserted the T_b_ thermistor 4-6 mm into the cloaca and attached its cable with liquid silicone to the feathers around the cloaca (Ricklefs & Williams 2003). Simultaneously, we measured T_a_ every 10 seconds with a second thermistor at the bottom of each chamber. For 15 out of 249 trials, the recording of T_a_ was lost, so to these we assigned the median of the median T_a_ s from the same night. Thermistors were inserted into the chamber through a lateral hole that was sealed with silicone (Fig. S2). Prior to deployment, we calibrated thermistors at cold (-4 °C) and warm temperatures (25 °C) against a laboratory-grade digital thermometer. When hummingbirds did not endogenously arouse from torpor before 07:00 (N = 11), the trial was omitted, and we warmed birds passively and fed them (as we did with all other individuals) before release. We marked individual birds by cutting a small portion of a tail feather and performed only one trial per bird. Capture and handling procedures were approved by the Institutional Animal Care and Use Committees at the Icesi University and collecting permits we granted by National Natural Parks (# Resolución 2014, IDB0325).

*Defining torpor*

First, we qualitatively identified torpor from a continuous drop of T_b_ temperature down to a low level (generally near T_a_) where T_b_ remained relatively stable, until a spontaneous increase in T_b_ began that persisted until euthermic levels were reached (Fig. S3). For birds this increase in T_b_ is typically accomplished using endogenous heat production (Wang 1989; Calder 1994; Geiser 2004). However, for the purposes of this study, torpor is considered distinct from a hypothermic state of reduced T_b_ where spontaneous arousal is not observed (Tomlinson *et al.* 2007). In eleven trials birds were not observed initiating endogenous arousal from torpor before ca. 07:00; these cases did not allow hypothermia to be distinguished from torpor and were excluded from analyses of the usage of torpor. Moreover, 29 trials showed periods of incomplete data acquisition due to sensor failure during the night or were not unanimously classified as torpor use by all three torpor definitions (see below). Because a bird may have used torpor when the sensor failed and we would not have been able to detect it, these trials too were excluded.

Although torpor exists along a continuum (Mckechnie & Lovegrove 2002; Schleucher 2004; Ruf & Geiser 2015; Shankar *et al.* 2022) practitioners often choose from a variety of similar, yet somewhat arbitrary, thresholds. For example, some authors defined torpor bouts using an absolute T_b_ threshold (if T_b_ ever drops below 30º; C e.g. Brigham *et al.* 2000; Wolf *et al.* 2020), while others have used relative T_b_ thresholds (drop in T_b_ by 5 º C below euthermic resting temperature (Brigham 1992; Ruf & Geiser 2015). We implemented both torpor definitions (defining resting temperature as the median euthermic T_b_ before midnight 00:00 or before the obvious descent into torpor, whichever came first; see thick horizontal lines in Fig. S3A, S3C). We also calculated metrics that were useful in qualitatively separating torpid birds' temperature curves from those of normothermic birds: for each trial, we determined the distribution of the temperature gradient between the T_b_ and the median T_a_ (which remained largely stationary within a night). From this distribution of differences, we calculated the mean and variance (Fig. 3B, S3D). Torpid birds had lower means and higher variances of the distribution of the differences between T_a_ and T_b_ than normothermic birds (Fig. S4). All torpor definitions agreed in over 94% of all trials. While the two threshold-based definitions agreed in over 94% of all trials, we also classified each trial manually, looking for characteristic drops in T_b_, and using the mean and variance of the frequency distribution of differences between T_a_ and T_b_ (Figs. S3-4). We used this manual definition of torpor for subsequent analyses, as it showed >95% agreement with both absolute and relative threshold-based torpor definitions.

*Characteristics of torpor*

We calculated torpor bout duration (TBD) in three complementary ways (Fig. S5). For trials with complete data, we defined complete TBD as the duration from the beginning of the entry into torpor until the end of the exit from torpor (orange, N = 63, Fig. S5A). Because in some trials, data recording ended before the return to normothermic euthermy was completed, we also defined short TBD as the duration from the beginning of the entry into torpor until the beginning of the exit from torpor (red, N = 78, Fig. S5AB). Finally, we defined incomplete TBD as including trials with birds that clearly used torpor to drop their Tb, although either the beginning of entry into torpor or the beginning of the exit from torpor were not recorded (yellow, N = 93, Fig. S5A-C). Incomplete TBD has the highest sample size but underestimates TBD, and complete TBD appropriately describes the full departure from normothermy but has the lowest sample sizes. To balance data quantity and quality, we emphasized the results of the short TBD, which has intermediate sample sizes and only fails to capture the exit from torpor, which was generally short. To be additionally cautious, we only show results in the main text that were consistent across short TBD and at least one other TBD definition (Table S2).

Upon classifying a bird as employing torpor, we determined three timepoints of interest for each trial using a custom interactive data visualization Shiny application in R software (R Development Core Team 2012; Chang *et al.* 2021). These were: 1) the beginning of the entry into torpor, 2) the beginning of the exit from torpor and 3) the end of exit from torpor. The Shiny application visualized the raw data and used the empirical first differences in subsequent temperature measurements (analogous to a first derivative) to reveal turning points in temperature curves (where first differences crossed 0). We located the beginning of the entry into torpor as the point in time preceding a sustained descent in T_b_ that began at euthermic levels and dropped precipitously toward T_a_. At this point, the first differences departed from 0 and remained negative. We located the beginning of the exit from torpor as the point in time at which T_b_ began a rapid, sustained ascent from near T_a_ to euthermy. At this point, the first differences departed from 0 and remained positive (Fig. S6, left side of selected temperatures in blue). We located the end of the exit from torpor as the point in time at which T_b_ returned to euthermy. At this point, the first differences dropped back down to 0 (Fig. S6, right side of selected temperatures in blue). After selecting the beginning and end of the exit from torpor, we defined the rate of the exit from torpor as the slope of ln(T_b_) over ln(exit duration (minutes)). This is shown in the bottom right panel of Figure S6.

*Body condition*

Body condition in birds is usually assessed by measuring fat scores (Krementz & Pendleton 1990) or by extracting residuals from the regression of body mass on a phenotypic measure of body size (e.g. tarsus length). In passerines, body size is preferably estimated by tarsus length (Rising & Somers 1989; Senar & Pascual 1997), as mean body mass and tarsus length are highly correlated across species (Pearson’s correlation coefficient ρ = 0.817, data from Tobias *et al.* (2022) and Wilman *et al.*, (2014)). However, hummingbirds live aerial lifestyles and tarsus length is likely constrained. Compared to passerines, average hummingbird tarsus length is 38% less strongly correlated with body mass across species (ρ = 0.589, Fig. S7A), so it might not be useful for calculating body condition. Moreover, the average hummingbird intraspecific standard deviation of tarsus length (0.631, data from Tobias *et al.* (2022)) is less than half of the average passerine intraspecific standard deviation of tarsus length (1.239), further underscoring the constraints on hummingbird tarsi (Fig. S7B). Thus, it is unclear to what degree tarsus length is an appropriate phenotypic measurement of body size in Trochilidae.

We lacked morphometric measurements of body size for the birds in our study of torpor so we turned to an additional data set we have collected from ongoing mistnetting efforts (N = 1,035 cleaned records of adults of focal species collected over six years) to explore morphological measurements of body size and to calibrate proxy variables of body condition. We found that tarsus length was uncorrelated with body mass within each species and correlated with body mass only across species (Fig S8A, D). Bill length was uncorrelated with body mass within all species but one and was correlated across species (Fig. S8B, E). Wing length was correlated with body mass within ten out of 21 species and was correlated with body mass across species (Fig. S8C, F). We exploited this weak intraspecific allometry between body mass and morphometric measures of body size to define a proxy variable for body condition that we could calculate from the birds used in the torpor trials that lacked morphometric measurements: this was the difference between the individual’s initial body mass and the species’ average body mass. Given that the allometry between body size and body mass is generally weak within the species in our study, we reasoned the individual differences from species-specific average adult body mass should approximate a measure of body condition that accounted for species-specific size variation.

We first calibrated this definition of body condition against a size-corrected definition of body condition using the extended mist-netting data from 1,035 individuals of the 29 species in our study. The residuals from models of species-specific body mass given known tarsus length, bill length, or wing length were highly correlated with our proxy definition of body condition across species (Fig. S9, all P < 0.001). As all measures of body condition were highly correlated with each other across (Fig. S9) and within each well-sampled species (Figs. S10-12), we believe that for the purposes of our study, we are justified in defining body condition in the birds from our torpor study as the difference in body mass to species-specific averages from all our body mass data (Fig. S13). Nevertheless, it is worth noting that although this proxy variable of body condition appears appropriate in the species of our study (at least as a candidate predictor variable of characteristics of torpor), it remains only a proxy variable of body condition. Other measurements (e.g. fat scores, muscle volume, crop volume, other phenotypic measurements) might turn out to be better indicators of body condition in hummingbirds.

*Sustained temperature gradient*

By calculating the sustained temperature gradient (STG) (Fig. S14), we were able to quantify variation in torpor depth along its continuum from deep to shallow torpor (Mckechnie & Lovegrove 2002; Schleucher 2004; Ruf & Geiser 2015; Shankar *et al.* 2022). Determining the STG over the entire night was straightforward for torpid and normothermic birds (Fig. S14C), but calculating the STG from the beginning of the night until the beginning of the exit from torpor was only possible for torpid birds (Fig. S14B). Even though normothermic birds did not enter torpor, we calculated their pre-torpor STG by setting t_1_ equal to the first time of T_b_ recording and setting t_2_ as equal to the average time of beginning entry into torpor at that station (Fig. S14A). At Anchicayá birds initiated the descent into torpor on average at 00:42 (range: 21:42 – 03:42); at Chicoral birds entered torpor on average at 23:42 (range: 21:24 – 02:06); and at Farallones, birds entered into torpor on average at 23:30 (range: 20:48 – 02:12). We consider this approach appropriate as it lets us capture the variation in STGs that occurs for all birds in the general time period before the obvious torpor-defining drop in T_b_.

Although STGs over the whole night were significantly correlated with nightly loss of body-mass (t = 4.297, df = 180, P < 0.0001, ρ = 0.305), inferring energy expenditure from temperature data alone is challenging; we therefore considered three major issues that might mask the signal of energy expenditure in T_b_. First, we acknowledge that small-scale variation in metabolic rate (MR) contributes to energy expenditure. It has been noted that instantaneous decreases in MR during torpor can precede the apparent entry into torpor that would be discerned from T_b_ alone by some minutes (M. Chappell, F. Geiser personal communication). Thus, variation in T_b_ alone might not always be proportional to MR when sampled at high temporal resolution. However, by integrating differences in T_a_-T_b_ gradients over long portions of the night (and even the entire night) we deliberately decreased the temporal resolution of measurement and thus suspect that this concern might be at least partially alleviated.

Second, variation in energy expenditure can also be driven by variation in thermal conductance. This is important, as conductance varies across species: birds with low thermal conductance inhabit cold areas of the globe (Fristoe *et al.* 2015). Moreover, conductance can be modified within species by behaviors such as group-huddling (Gilbert *et al.* 2008, 2010) and by modifying plumage thickness (Wolf & Walsberg 2000). Although we cannot address the potential for intraspecific, behavioral modification of conductance in our study, we can evaluate the potential for interspecific differences in conductance. Therefore, we collected data on conductance and body mass from 23 species of hummingbirds from the literature (Fristoe *et al.* 2015; Londoño *et al.* 2017). Interspecific variation in conductance in hummingbirds is highly predicted by body mass in a phylogenetic regression (N = 23, ln(Body Mass) = 0.616, SE = 0.143, T = 4.31 , p = 0.0003, R^2^ = 0.469; Fig S15A) and shows low phylogenetic signal (λ = 0.002). Thus, we suspect that at least some portion of interspecific variation in conductance (which might be present in our dataset of 29 species for which we measured torpor) can be accounted for by body mass, which we consider as a potential explanatory variable in our analyses.

Third, variation in energy expenditure can also be driven by variation in T_lc_, as the width of the thermoneutral zone can be adjusted by raising or lowering T_lc_. Although we cannot evaluate the potential for intraspecific variation in T_lc_, we consulted literature estimates of avian T_lc_ (Fristoe *et al.* 2015; Londoño *et al.* 2017). Only five of our species had T_lc_ estimates available from the literature and only 19 estimates of T_lc_ from other Trochilidae were available, so we imputed the missing T_lc_ data from a global dataset of T_lc_ values of birds (N = 453). We think that imputation in this case is justified because body mass had a significant effect on T_lc_ in birds in a phylogenetic regression of T_lc_ on body mass (ln(Body Mass) = -2.095, SE = 0.345, T =-6.07 , p < 0.0001, R^2^ = 0.078, Fig. S15B). Because this regression showed a strong role of phylogeny (λ = 0.774), we took phylogenetic correlations into account while imputing the missing T_lc_ from the species in our study (Garland & Ives 2000). As observed nocturnal T_a_ in our experiments (mean = 19.02 °C, SE = 4.44, range: 10.91 – 26.79 °C) were consistently below imputed T_lc_ (mean = 29.98 °C, SE = 0.75, range: 28.63 - 31.60 °C), we can assume that interspecific variation in T_lc_ is not a main driver of T_b_ dynamics in our study. For all 235 trials in which we had the ability to diagnose torpor, all but two experienced median T_a_ levels below the species-level T_lc_. These two trials occurred for *Doryfera ludovicae*, which once entered torpor and once remained in normothermy (with no effect of torpor definition). Thus, we have no reason to believe that T_a_ must reach extremely far below T_lc_ to initiate torpor. In any case, for all species in our study, nocturnal T_a_ appeared sufficiently low (beneath or close enough to observed and estimated T_lc_) to trigger a response in T_b_; therefore we suspect that variation in energy expenditure that is attributable to T_lc_ may be negligible for the purposes of our study.

The comparative data on T_lc_ and thermal conductance were analyzed with phylogenetic regression models in `phylolm` (Tung Ho & Ané 2014) using a summary tree from the entire Global Bird Phylogeny (Jetz *et al.* 2012).

*Phylogenetic hypotheses*

Our dataset consisted of multiple measurements of different species of hummingbirds, for which phylogenetic relationships are established (Jetz *et al.* 2012; McGuire *et al.* 2014). Thus, we accommodated both the phylogenetic non-independence of the data as well as the repeated measurements per species in our analyses and used a summary tree (maximum clade credibility) from a recent version of the Global Bird Phylogeny (Jetz *et al.* 2012) with the Hackett backbone (Hackett *et al.* 2008), as this tree corresponded closely to a summary tree from another phylogenetic study (McGuire *et al.* 2014) and had preferable taxonomic coverage. Our dataset included one taxon (*Phaethornis striigularis*, the Stripe-throated Hermit) that is not present in the phylogenetic tree provided by (McGuire *et al.* 2014) but present in the Global Bird Phylogeny (Jetz *et al.* 2012). Therefore, we preferred the Global Bird Phylogeny to maximize species coverage. Both phylogenies showed widespread agreement (Fig. S16). For analyses involving species beyond our analytic dataset, we used a summary tree from the Global Bird Phylogeny following Baldwin *et al.* (2022).

*Statistical analysis – torpor experiments*

We investigated the drivers of torpor, variation in its characteristics, and consequences of torpor with Bayesian phylogenetic generalized linear mixed models (PGLMM) implemented in the brms package (Bürkner 2017, 2018; Carpenter *et al.* 2017). This approach allowed us to simultaneously incorporate phylogenetic non-independence, repeated sampling of species, and non-Normal response distributions (de Villemereuil & Nakagawa 2014).

We used phylogenetic generalized linear mixed models (PGLMMs) to identify which fixed effects best explained variation in the response variables. We structured the PGLMMs to share intercepts according to their phylogenetic relatedness and, additionally, fitted a random effect term for the repeated measurement of each species. For all questions, we fitted a series of regression models and began model selection with a full model that included all additive single Z-transformed predictor variables as well as all two-way interactions between the remaining predictor variables. Model selection proceeded by eliminating each term one by one and refitting the simplified model. We simplified models by stepwise elimination of weak and uncertain predictors (eliminating predictors whose posterior medians were close to zero and whose 95% credible intervals (CI) overlapped widely with zero) while ensuring model nestedness (i.e. an additive effect cannot be eliminated when it is still involved in a significant interaction term), until significant predictors remained. We considered coefficients to be significantly different from zero if their 95% CIs did not overlap with zero. If the 95% CI did, but the 90% CI did not overlap with zero, we considered the coefficient to be marginally significant. We report all model coefficients from the final time they featured in model selection, or from the final model (Tables S1-S4), and also provide supplemental figures that illustrate the effect sizes during model selection procedure (Figs. S19-S28).

Because the sustained temperature gradient contained up to 57 missing values, we first performed model selection without this variable (its inclusion reduced the sample size considerably). Upon obtaining a final model, we added the sustained temperature gradient (and its interactions with the remaining variables) back into the model and continued model selection on the reduced dataset.

All models were fitted in the brms package using stan (Bürkner 2017, 2018; Carpenter *et al.* 2017). We fitted each model with four chains with 25,000 iterations, a warm-up period of 15,000 iterations, and a thinning rate of 10. We assessed convergence by manually inspecting traceplots and ensuring that each parameter’s Gelman-Rubin statistic fell below the threshold value of 1.1 and ESS values were uniformly high (well above 1,000). To avoid predictor collinearity, we confirmed that variance inflation factors were below <10 on full and final models (Dormann *et al.* 2013).

*Statistical analysis – torpor experiments with expanded dataset*

We also considered the effect of adding the raw data from a recent report of torpor in Andean hummingbirds at higher elevations (3,800m asl; Wolf et al. 2020), for which we also calculated the pre-torpor STG as previously. However, these data did not permit us to evaluate the effect of individual body mass or body condition as specimen-level data were not provided; therefore, we queried species mean body mass from the database “Elton Traits” (Wilman *et al.* 2014). Nevertheless, we were able to perform our calculation of pre-torpor STG to determine the variables that drive the usage of torpor (Fig. S20).

*Statistical analysis – phylogenetic propensity for torpor and phylogenetic signal*

We tested for phylogenetic signal in the PGLMMs by calculating the proportion of random effect variance attributable to the phylogenetically pooled random intercepts, following the brms vignette (<https://cran.r-project.org/web/packages/brms/vignettes/brms_phylogenetics.html>; Table S5). This quantity corresponds to Pagel’s λ in most cases (c.f. Barrow *et al.* 2019; Wolf *et al.* 2020), although it has not been explicitly extended to non-Gaussian response distributions. We also extracted the phylogenetic random effect levels from the final model of each response; these levels are the species-specific deviations from the predicted average response, after taking into account all other model variables. To these species-specific values, we also applied the function ‘phylosig’ from the phytools package (Revell 2012) to calculate Blomberg’s K. We tested for phylogenetic signal using 1,000 simulations, and also used the test for Blomberg’s K with trait observation errors that we defined as the posterior standard error of the species-specific random intercepts. Finally, we also calculated Pagel’s λ in the R package motmot (Thomas & Freckleton 2012), using two chains and default runtime conditions (20,000 iterations, 10,000 burn-in, thin rate = 10) and this achieved satisfactory convergence (Table S6).

*Elevational ranges*

We obtained data on elevational ranges from a recent database (Quintero & Jetz 2018) that provided each species’ Andean elevational range within a country. We averaged each species’ maximum and minimum elevational ranges across countries to obtain average elevational Andean range limits (Fig. 5). We calculated the elevational midpoint as the midpoint between these average maxima and minima.

*Statistical analysis – evolutionary models*

We first fitted a linear model to describe the relationship between elevational midpoint and species-specific propensity for torpor. We then refitted the linear model as a phylogenetic regression model (PGLS) using Pagel’s λ. Finally we used the package sensiPhy (Paterno *et al.* 2018) to address phylogenetic uncertainty in the PGLS by refitting it on a sample of 100 randomly selected trees from the posterior tree distribution with the Hackett backbone (Hackett *et al.* 2008; Jetz *et al.* 2012).

As phylogenetic correlation does not imply causation, we turned to a recent process-based evolutionary model to test the effect of changes in propensity for torpor on the rate at which changes in elevational midpoint evolve (Hansen *et al.* 2021). Both variables were qualitatively Normally distributed and were first transformed to become positive. As recommended (Hansen *et al.* 2021), we ensured that changes in elevational midpoint in our sample (N = 29) could be appropriately modelled by Brownian motion (BM), using the fitContinuous function in the geiger package (Pennell *et al.* 2014). We obtained estimates of evolutionary rate parameters (σ^2^ = 0.021) in this exploratory model. We then used the “BM1” model from Hansen *et al.* (2021) to test the following hypothesis: changes in torpor propensity influence the rate of evolution for species elevational midpoints, given a BM model with a σ^2^_elevation_ that is dependent on the propensity for torpor. Given that we were focusing on a group of taxa that have diverged over tens of millions of years, we believe that the pattern of elevational evolution is primarily macroevolutionary; therefore, we selected the “BM1” model, as suggested (Hansen *et al.* 2021). We then calculated the rate-regression parameters *a* and *b*, and bootstrapped the model to obtain uncertainty estimates in the evolvability package (Bolstad *et al.* 2014; Hansen *et al.* 2021).

Next, as our torpor data represented only a small portion of the hundreds of Andean hummingbirds, we expanded the scope of our study and explored patterns of elevational variation across 206 hummingbird species with data from Quintero & Jetz (2018) and macroevolutionary models in bayou (Uyeda & Harmon 2014), which implements Bayesian reversible jump Markov chain Monte Carlo (rjMCMC) models. We fitted a bayou model in which traits are assumed to follow a multi-optimum Ornstein-Uhlenbeck (OU) model of trait evolution, although shifts between optimum trait value are allowed. In this sense, we used bayou to reveal, locate and quantify shifts in elevation across Andean hummingbirds in order to establish where and when shifts in elevation have occurred. In contrast to the previous model of evolutionary rates (Hansen *et al.* 2021), the bayou model assumes that elevational midpoints evolve as a multi-optimum OU process. We believe this modelling approach is justified for two reasons: first, with the reduced sample size above (N = 29), we lacked the statistical power to reject the null hypothesis of the parsimonious Brownian motion model. Second, when viewed across numerous Andean species, the composition of hummingbird communities shows strong phylogenetic structure across elevations (Graham *et al.* 2009), with different clades often replacing one another across elevational zones (Stiles 2004; McGuire *et al.* 2014): this pattern suggests that a multi-optimum OU model can be appropriate as it can permit different groups to be characterized by different elevational optima.

We used a uniform empirical prior on elevation (constraining it to sea-level and the maximum height of the Andes, 6,962 m asl, which is well beyond the highest elevational range of any hummingbird) and allowed only one shift per branch. We set a conditional Poisson prior on the number of shifts with a mean equal to 2.5% the total number of branches in the tree (206) and a maximum number of shifts equal to 5% (λ = 5.15, K_max_ = 10.3), following Uyeda *et al.* (2017). We ran three independent chains with 45,000,000 iterations each, used the first half as “burn in” and diagnosed convergence by visually inspecting traceplots and by verifying high effective sample sizes for all parameters (Table S7, Fig. S29). We retained only the shifts in elevation that were well-supported (posterior probability > 0.55) across all chains (Table S8, Fig. 5B).

Although we were using bayou in a deliberately exploratory manner and refrained from an explicit, model-comparison approach, we fitted a null model with shift numbers set to 0. We ran this model on a single chain for 10,000,000 iterations and failed to recover support for the same shifts in elevation as in the candidate model described above (Table S9). All analyses were conducted R Statistical Software, version 4.1.2. (R Development Core Team 2012).

**Supplementary Figures**

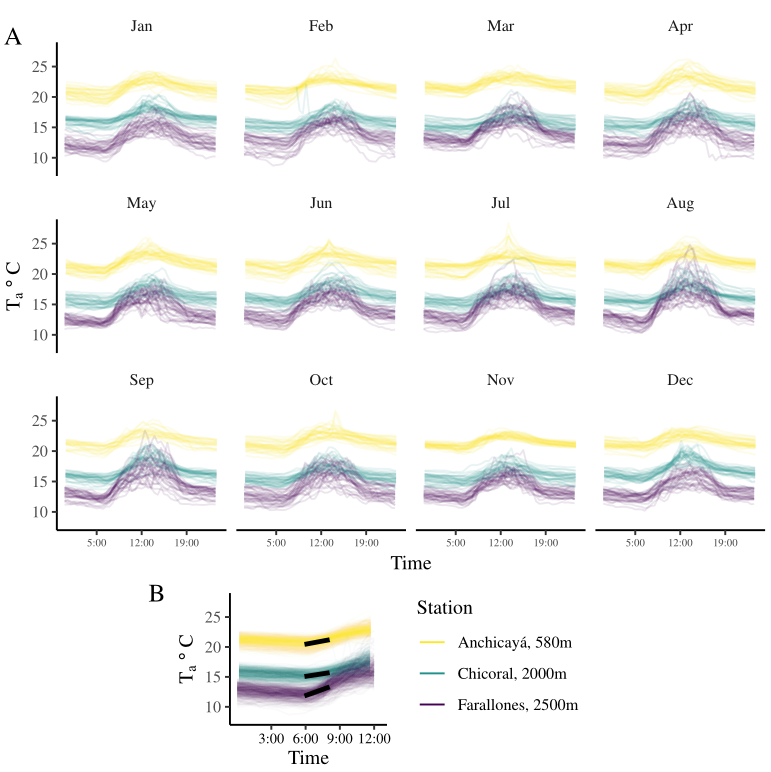

***Figure S1.*** *Environmental temperatures at field sites.* **A**) Circadian and circannual variation in ambient temperature along the elevational gradient. **B**) Circadian variation in ambient temperature along the elevational gradient in the mornings across all months. Each thin line shows T_a_ from one day of data collection between 2016 and 2019. Black lines show predicted T_a_ between 06:00 and 08:00: Slopes of time ± SE: Anchicayá (0.355 ± 0.037), Chicoral (0.291 ± 0.082), Farallones (0.688 ± 0.082).

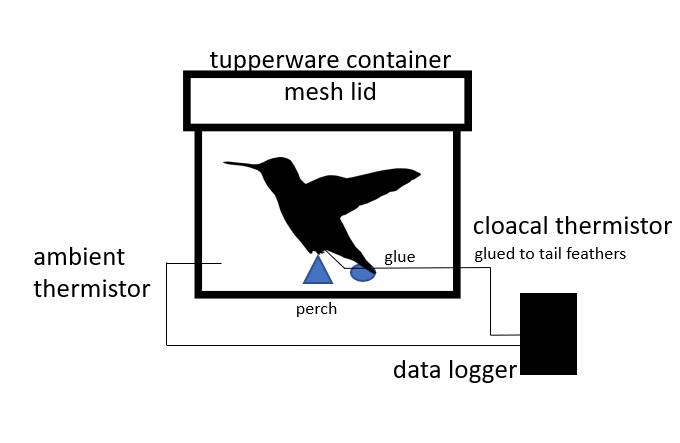

***Figure S2.*** *Experimental setup diagram.*

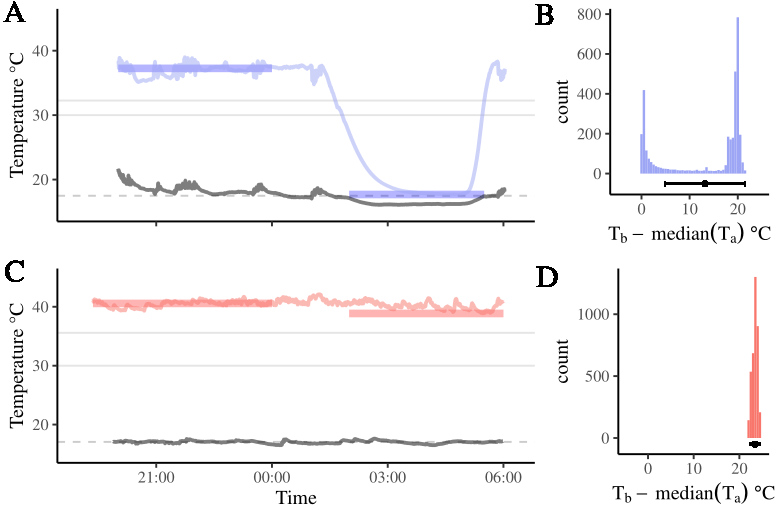

***Figure S3.*** *Typical temperature curves during torpor and normothermy*. Example of raw data in typical torpor (blue, **A, B**) and normothermic phenotypes (red, **C, D**). Ambient temperature shown in black points. In **A** and **C**, gray solid horizontal lines indicate the thresholds (30º C and 5º below resting T_b_). below which T_b_ would need to drop to identify torpor. Gray dashed horizontal lines indicate median T_a_. Thick red and blue lines indicate median resting T_b_ (left) and minimum T_b_ (right). Panels **B** and **D** show the distribution of differences between T_b_ and median T_a_. Horizontal black pointranges show the median and variance of the distributions of differences between T_b_ and median T_a_.

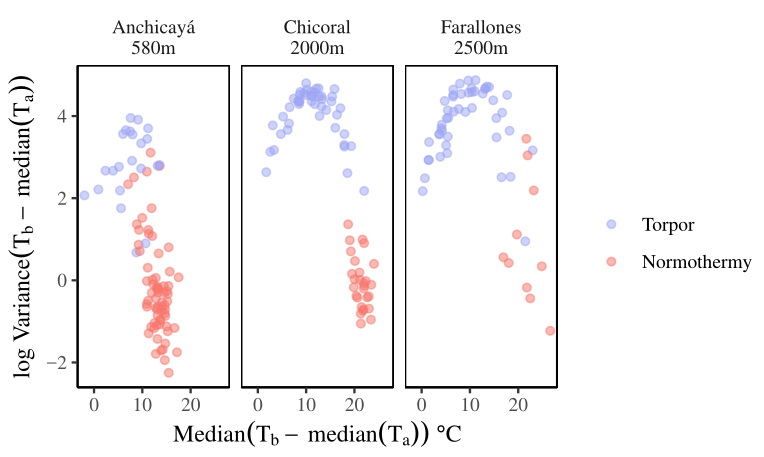

***Figure S4.*** *Across elevations, torpid birds were largely separatable from normothermic birds*: trials in which torpor was used had high variances of the distribution of differences between T_b_ and median T_a_. Normothermic trials showed low variances and high medians of the distribution of differences between T_b_ and median T_a_. Colors show manual torpor definition.

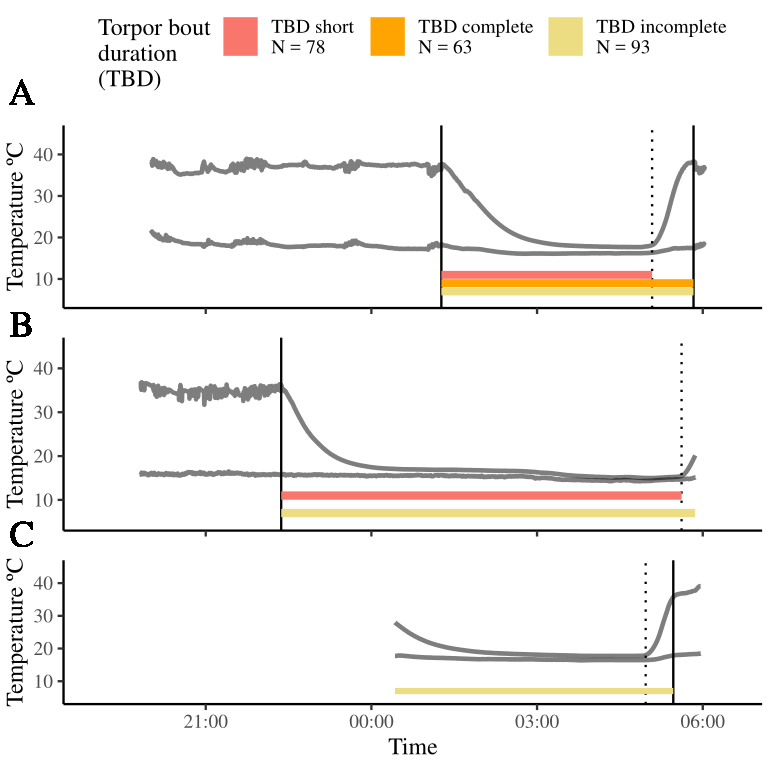

***Figure S5.*** *Definition of torpor bout durations.* We calculated three definitions of torpor bout duration (TBD). (**A**) Typical trial with all three TBD definitions available: complete TBD, short TBD and incomplete TBD. In trials like (**B**), data collection stopped before the completion of exit from torpor, permitting only the measurement of short TBD (red) and incomplete TBD (yellow). In other trials like (**C**), data collection failed to capture either the beginning of the entry into torpor or the beginning of exit from torpor, permitting only the measurement of incomplete TBD (yellow). Solid vertical lines indicate beginning of entry into torpor and end of exit from torpor. Dotted vertical lines indicate beginning of exit from torpor. Sample sizes for each TBD definition are shown in color legend.

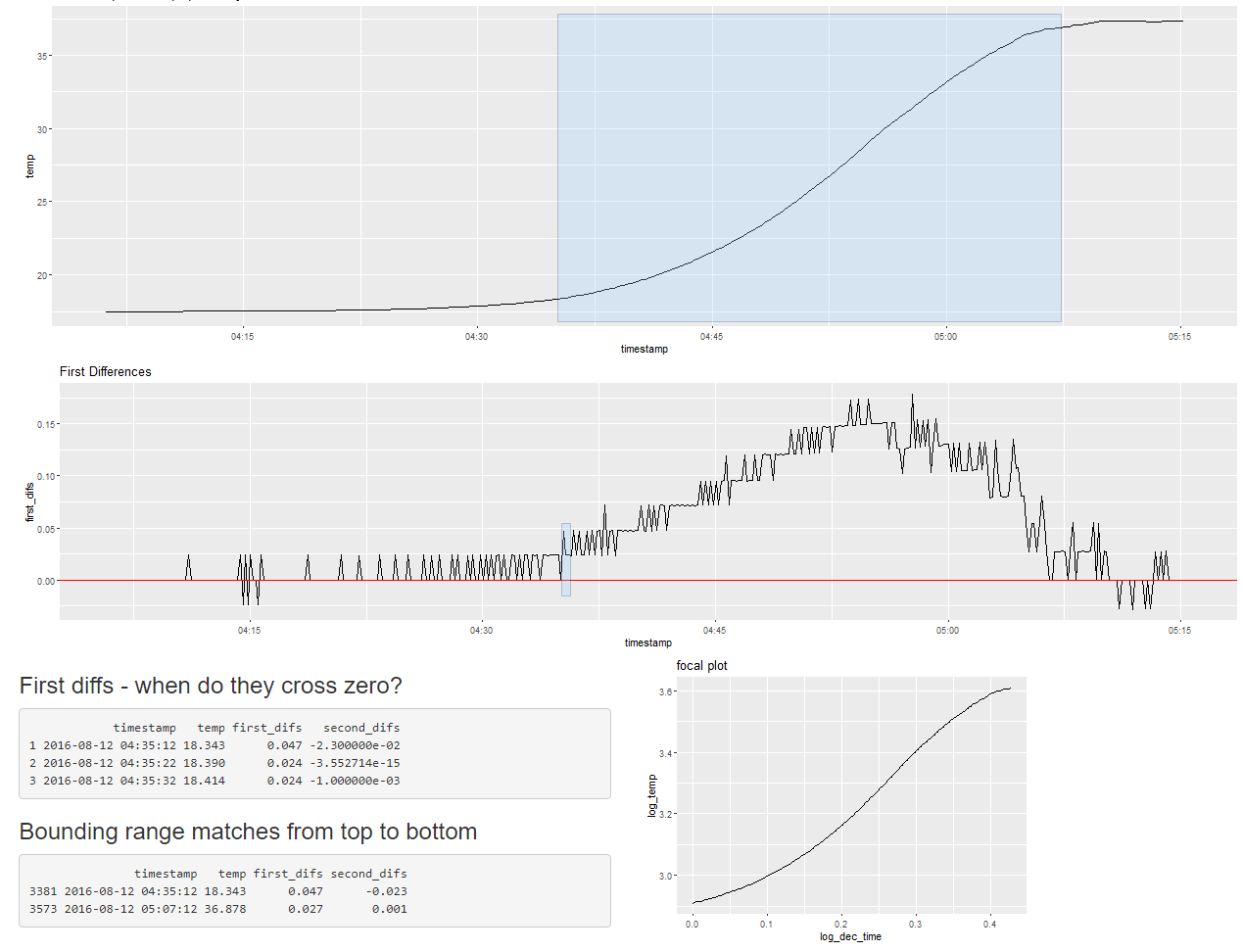

***Figure S6.*** *Screenshot of custom Shiny application in use - determining the time at which the exit from torpor began.* Top panel shows T_b_ through time, middle panels show first differences of T_b_ through time. By adjusting the selected portions of T_b_ (top panel), the middle panel and output in the tables in the lower left change. After determining the beginning and end of the exit from torpor, a log-log linear model is fitted to the data (bottom right panel) and the slope coefficient is returned. This application was used analogously to determine the rate of entry into torpor.

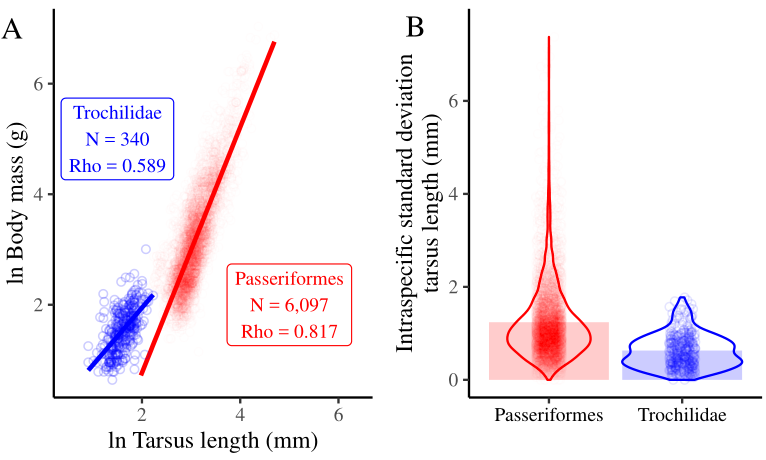

***Figure S7****. When quantifying body size, tarsus length is useful for passerines, but less useful for Trochilidae*. **A**) Across-species log-log linear models of mean body mass and mean tarsus length reveals different allometries and weaker correlation strengths in Trochilidae than in Passeriformes (data from Tilman *et al.* (2014) and Tobias *et al.* (2022). **B**) Average intraspecific variation in tarsus length is lower in Trochilidae than Passeriformes (data from Tobias *et al.* (2022).

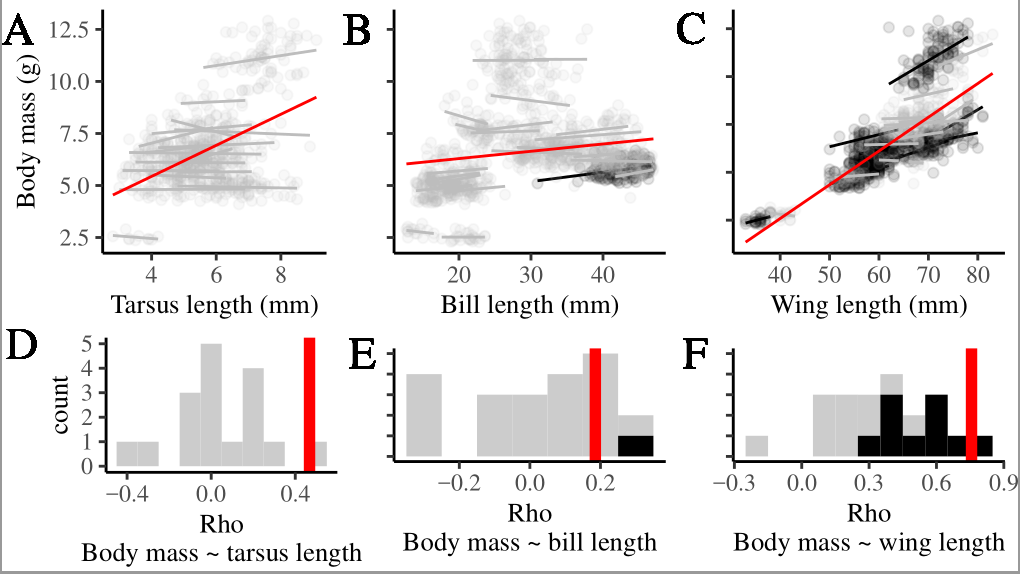

***Figure S8.*** *Body mass is highly correlated with body size across species but weakly correlated within species.* **A)** tarsus length, **B)** bill length, **C)** wing length. Data from external mist-netting efforts (N = 1,035 individuals of the species in our study). **A-C** Fitted lines are from linear models. Long red lines denote significant linear models fitted to all individuals; short lines denote linear models fitted to data for each species (black line: significant, gray: not significant). **D-F** show frequency distributions of intraspecific correlation coefficients (gray = non-significant, black = significant). Vertical red lines show significant correlation coefficients across all individuals.

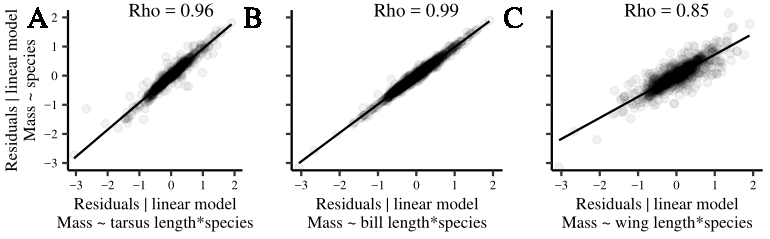

***Figure S9.*** *Calibration of an appropriate measure of body condition across species.* Body condition as defined by the deviation from species-specific average body mass (y axis; see Fig. S13) is highly correlated with body condition metrics that account for the allometric relationships (x axes) between **A**) tarsus length, **B**) bill length, or **C**) wing length with body mass (N = 1,035, p-values < 0.0001). Model formula shown in axis titles, where ~ denotes “as a function of” and * denotes the presence of additive effects and an interaction.

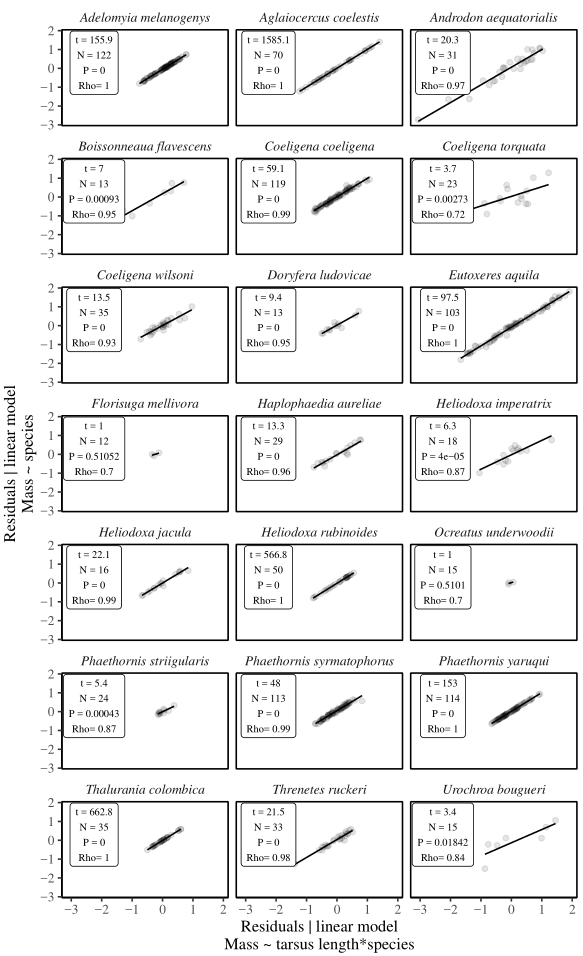

***Figure S10.*** *Calibration of an appropriate measure of body condition within species.* Body condition defined as the deviation from species-specific average body mass (y axis; see Fig. S13) is highly correlated with the body condition defined as residuals from the allometric relationship within species between tarsus length and body mass (x-axis). Model formula shown in axis titles, ~ denotes “as a function of” and * denotes the presence of additive effects and an interaction. Species with more than 10 occurrences in mistnetting data shown.

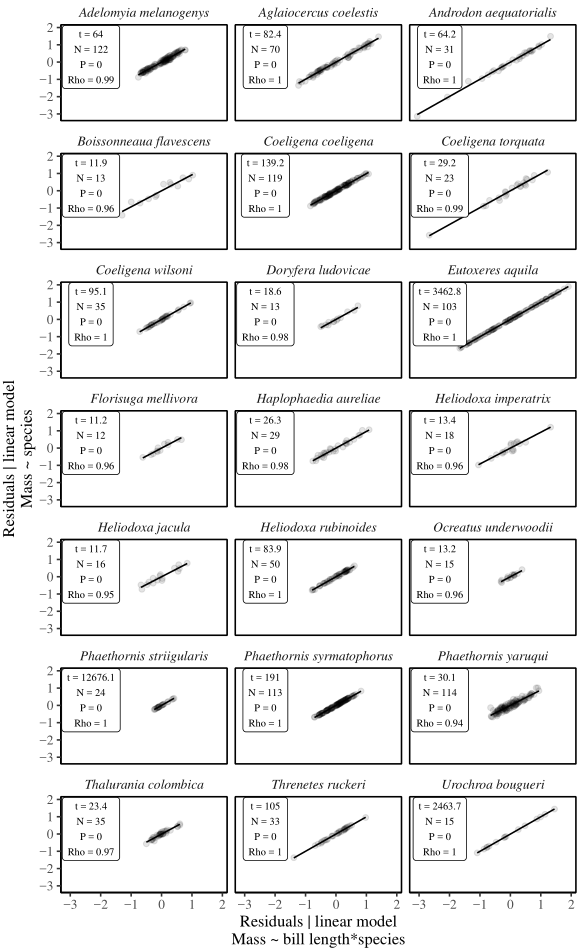

***Figure S11.*** *Calibration of an appropriate measure of body condition within species.* Body condition defined as the deviation from species-specific average body mass (y axis; see Fig. S13) is highly correlated with the body condition defined as residuals from the allometric relationship within species between bill length and body mass (x-axis). Model formula shown in axis titles, ~ denotes “as a function of” and * denotes the presence of additive effects and an interaction. Species with more than 10 occurrences in mistnetting data shown.

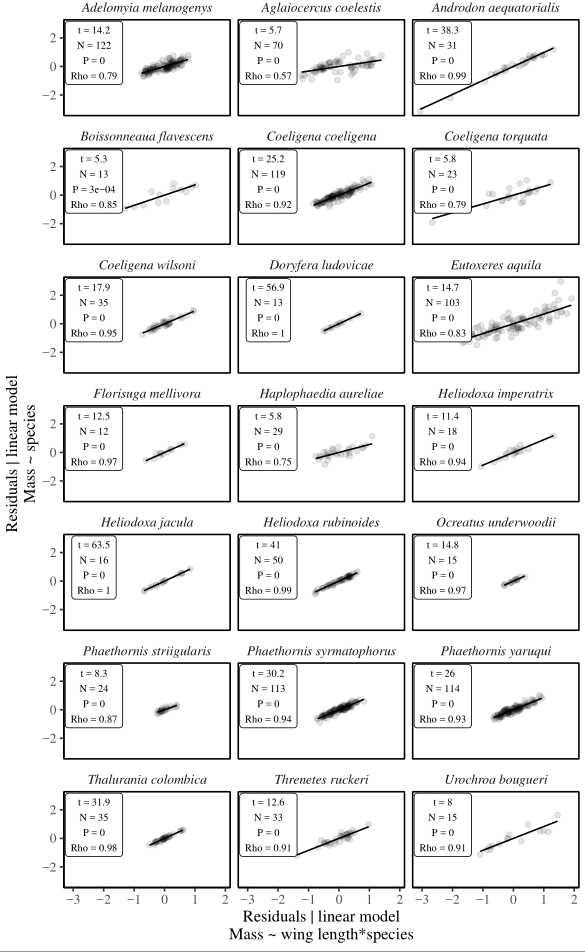

***Figure S12.*** *Calibration of an appropriate measure of body condition within species.* Body condition defined as the deviation from species-specific average body mass (y axis; see Fig. S13) is highly correlated with the body condition defined as residuals from the allometric relationship within species between wing length and body mass (x-axis). Model formula shown in axis titles, ~ denotes “as a function of” and * denotes the presence of additive effects and an interaction. Species with more than 10 occurrences in mistnetting data shown.

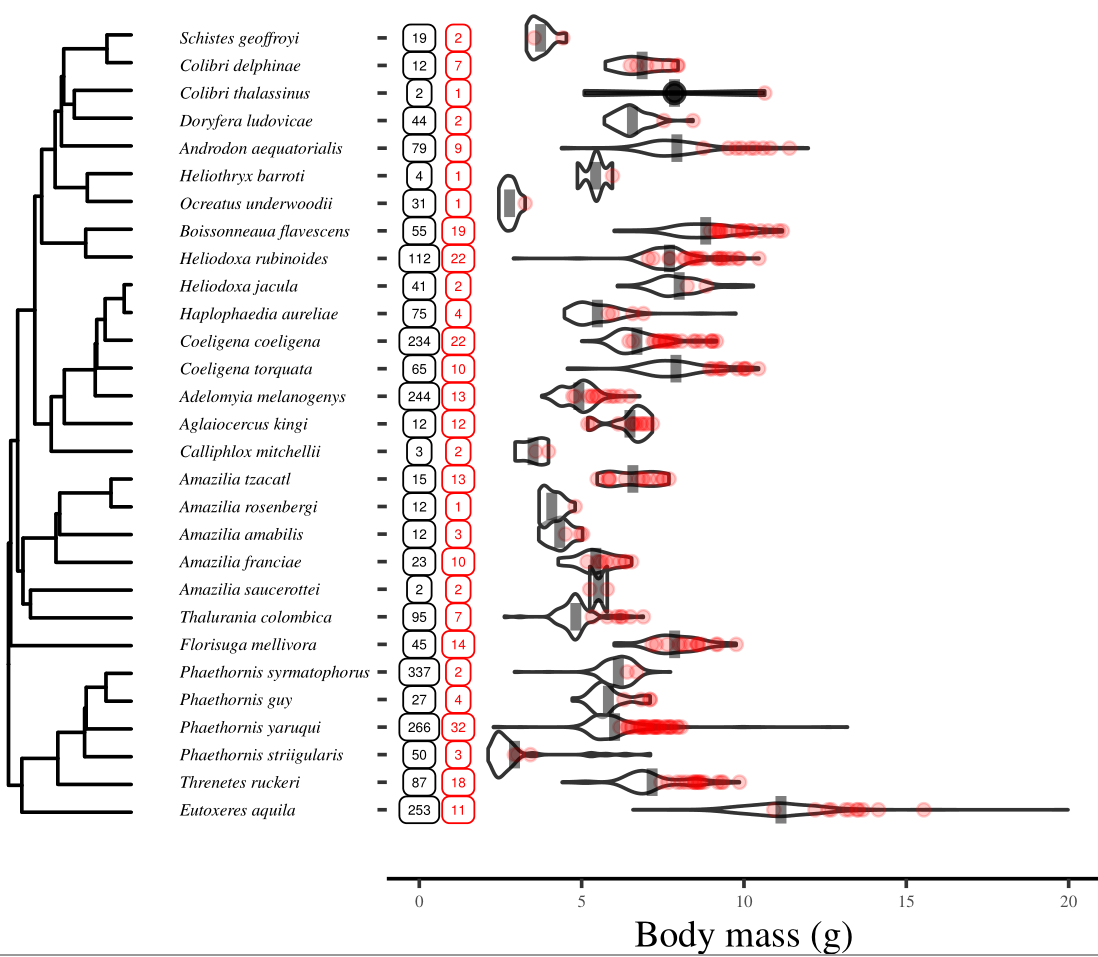

***Figure S13.*** *Body condition.* We defined body condition as the difference between individual body mass before torpor trials (red) and the species-specific average body mass from mist-netting data (black). Gray vertical line shows body mass averages, violin plots show densities, scattered points show body mass before torpor trials, and integers in squares show sample sizes (black: sample size in mist-netting data; red: sample size in torpor trials).

­­­Figure S14

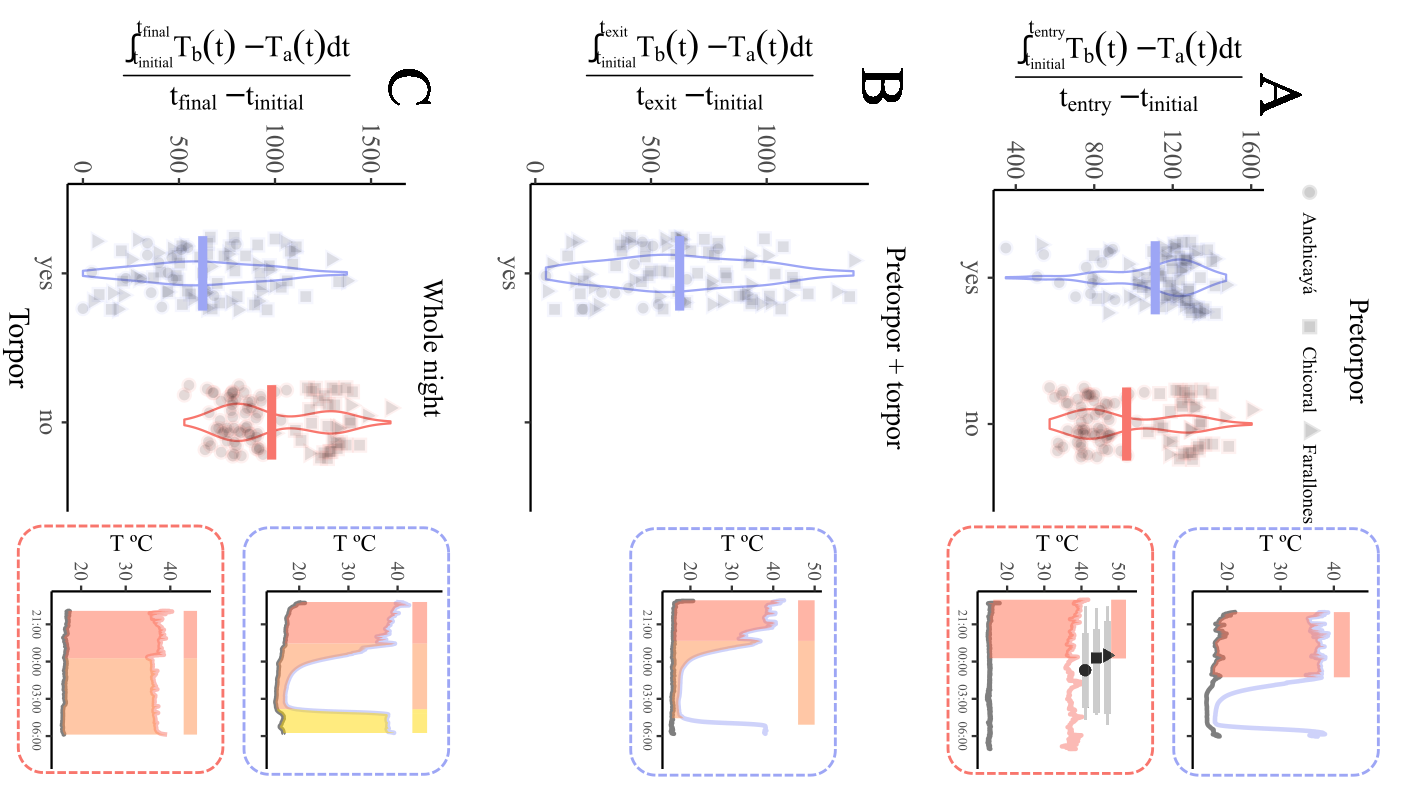

***Figure S14.*** *Defining the sustained temperature gradient (STG).* We calculated the sustained temperature gradient (STG) by integrating the difference between T_b_ (upper red and blue lines in right panels) and T_a_ (lower black lines in right panels) across each night. To standardize across recording durations, we divided this area by the duration of recording integration (temporal interval between t_1_ and t_2_). **A**) In the STG during the pre-torpor phase, the integration interval spanned the time of first recording and either the beginning of the entry into torpor (for torpid birds) or the station’s average time of the beginning of the entry into torpor (for normothermic birds – black shapes above normothermic T_b_ show mean, gray bars show standard error and range). **B**) In the STG during the pre-torpor and torpor phase, the integration interval spanned the time of first T_b_ recording and the beginning of the exit from torpor (for torpid birds only). **C**) In the STG for the whole night, the integration interval spanned the first and last times of T_b_ recording. In right panels, shading indicates the area between T_b_ and T_a_ and thick horizontal bars indicate durations (red: pre-torpor, orange: torpor, yellow: post-torpor). In left panels, points show STG values, violins show densities and horizontal bars show means.

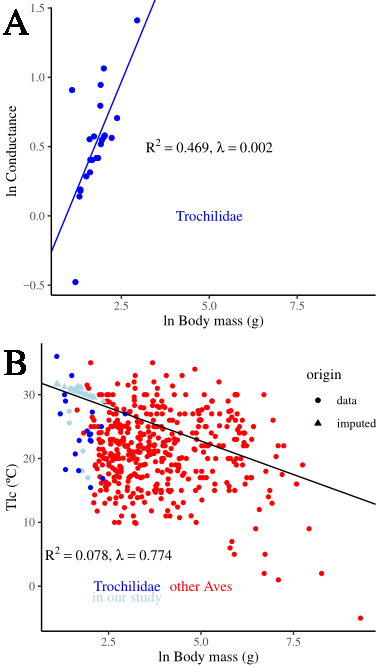

***Figure S15.*** *Interspecific variation in conductance and Tlc is linked to body mass.* **A**) In hummingbirds, conductance varies principally due to body mass in a sample of 23 species from the literature. Regression line is from a phylogenetic regression with Pagel’s λ using the consensus tree from the Global Bird Phylogeny. **B**) T_lc_ is significantly correlated with body mass in a sample of 437 species from the literature. The regression line is from a phylogenetic regression with Pagel’s λ using the consensus tree from the Global Bird Phylogeny and was used to impute the missing 24 T_lc_ data for our study species (light blue) following Garland and Ives (2000).

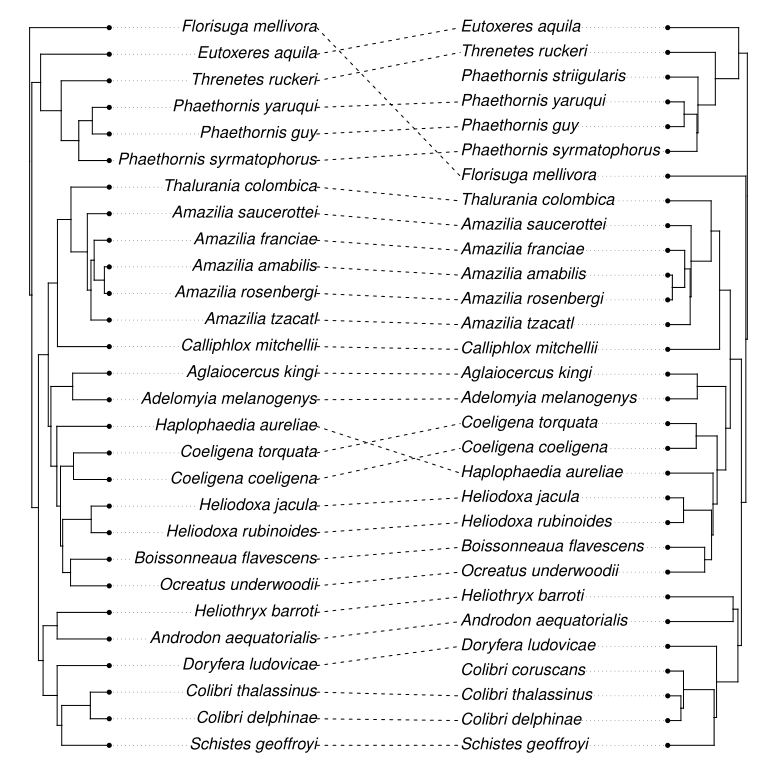

***Figure S16.*** *Comparison of phylogenetic hypotheses.* The phylogenetic hypotheses proposed by McGuire *et al.* (2014) (left) and the Global Bird Phylogeny (Jetz *et al.* 2012) (right) show widespread agreement.

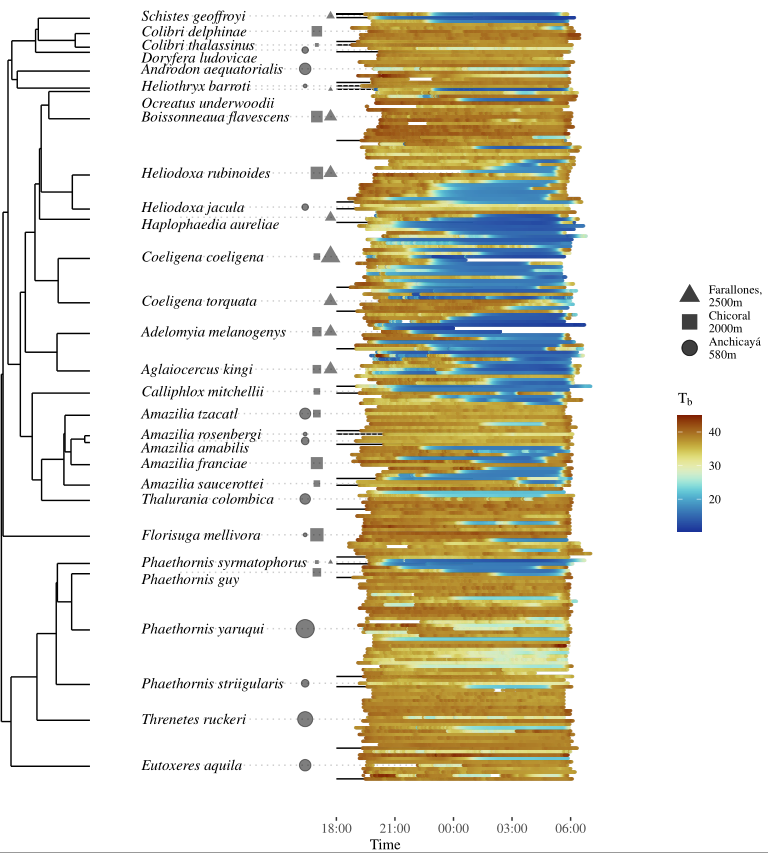

***Figure S17.*** *T_b_ of all individuals in phylogenetic order*. Torpor prevalence and depth is highly variable within and across species. All T_b_ data organized phylogenetically with the summary tree from the Global Bird Phylogeny from Jetz *et al.* (2012). Each horizontal colored line shows T_b_ of one nightly trial. Shape size is proportional to sample size at each station.

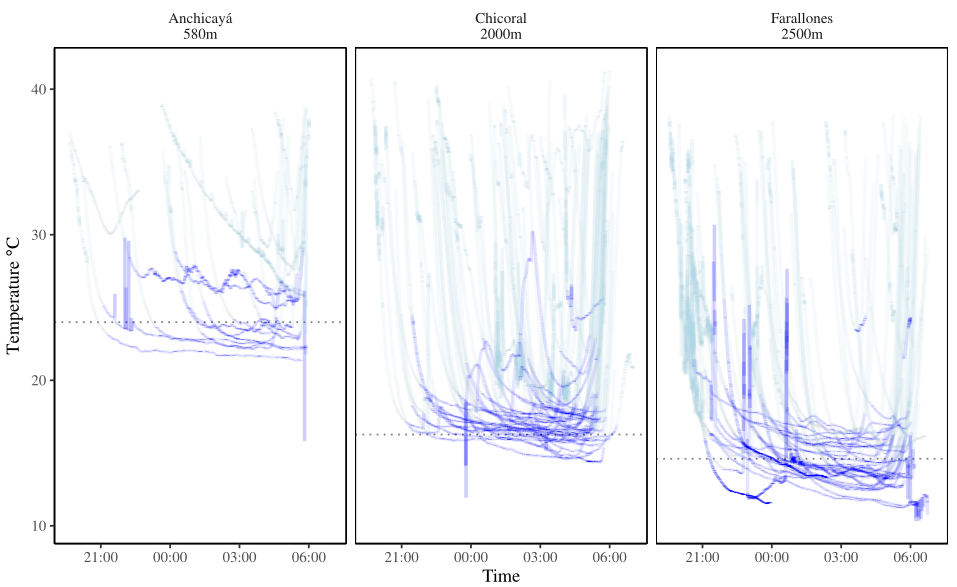

***Figure S18.*** *Torpor depth is highly variable across study sites.* Each line shows the measured Tb of a bird that entered torpor. Dark blue shows T_b_ after completing the descent into and before the ascent from torpor (each in light blue). Horizontal dotted lines show mean nightly T_a_.

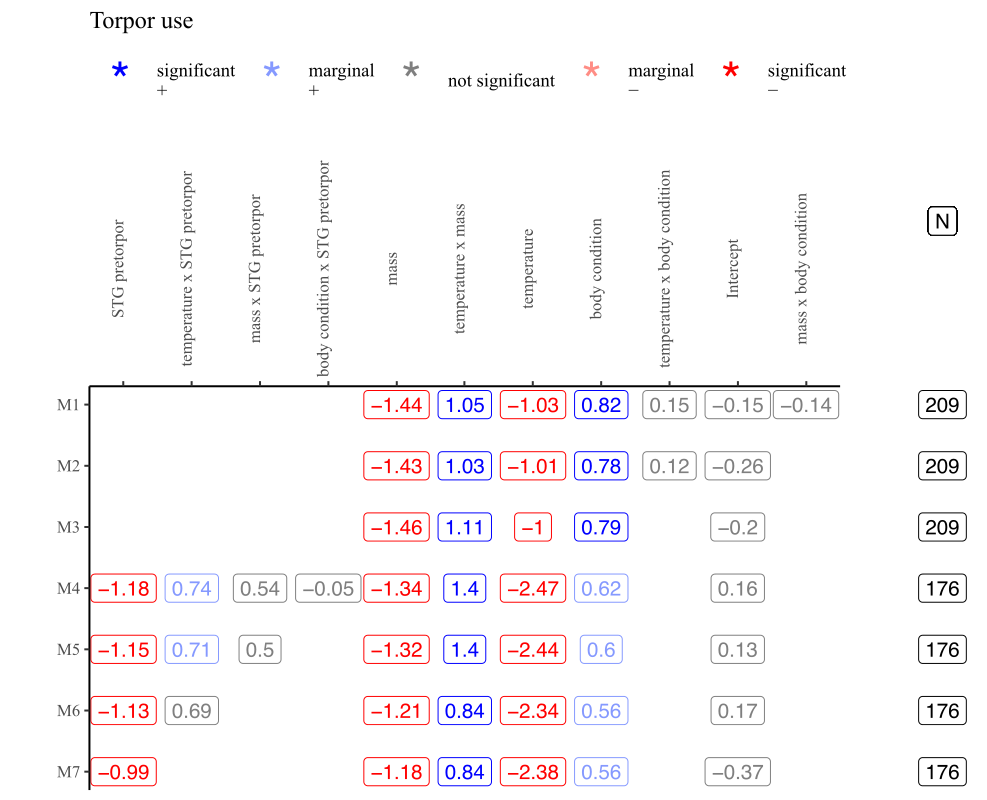

***Figure S19.*** *Graphical illustration of the model simplification process for variables that explained variation in the usage of torpor.* M1 represents the full model. N shows the variation in sample size due to considering the variable of pre-torpor STG.

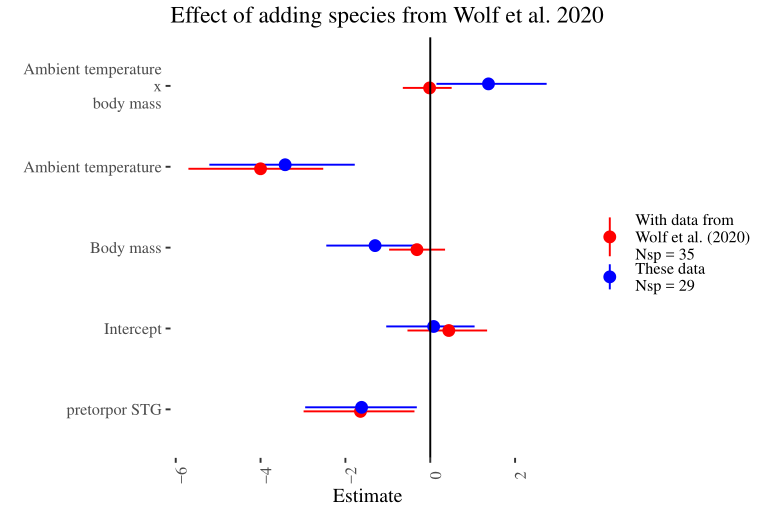

***Figure S20.*** *Adding the trials from Wolf et al. (2020) prohibited evaluation of the contribution of body condition and changed some of the inferred ecological drivers of the usage of torpor.* Coefficient point estimates are shown by points, and horizontal bars show width of the 95% credible interval.

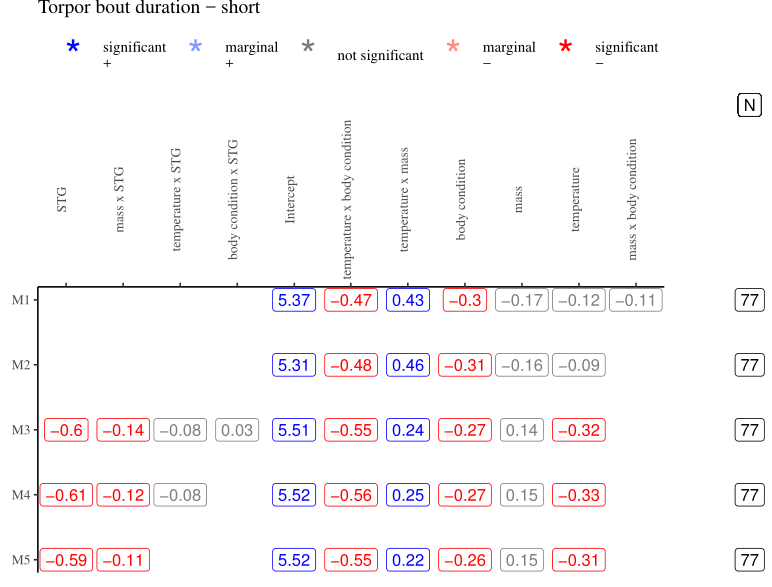

***Figure S21.*** *Graphical illustration of the model simplification process for variables that explained variation in short torpor bout duration.* M1 represents the full model. N shows the variation in sample size due to considering the variables of pre-torpor and torpor STG.

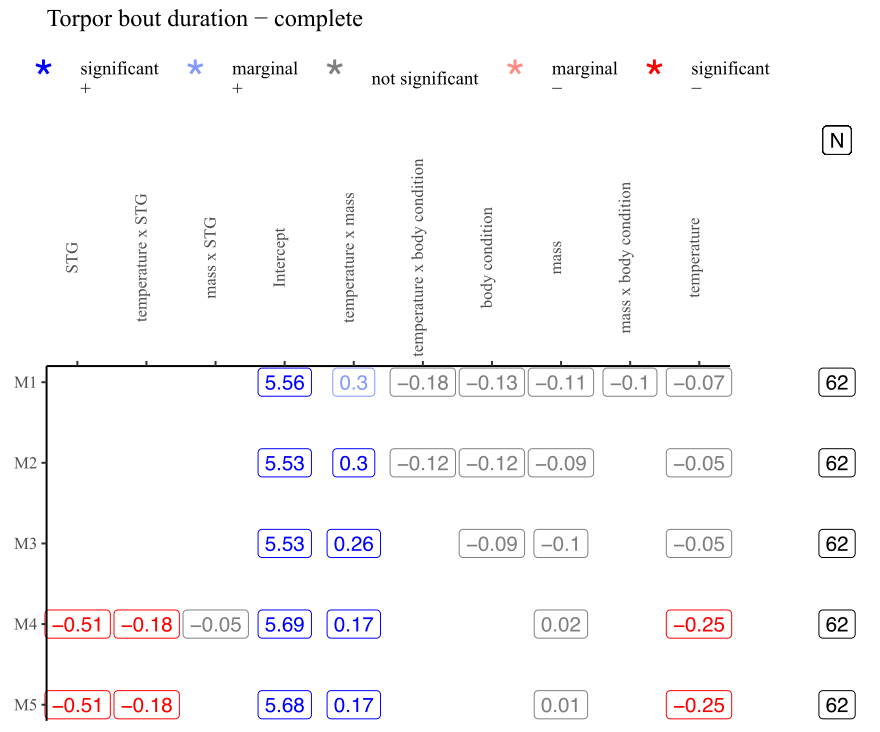

***Figure S22.*** *Graphical illustration of the model simplification process for variables that explained variation in complete torpor bout duration.* M1 represents the full model. N shows the variation in sample size due to considering the variables of pre-torpor and torpor STG.

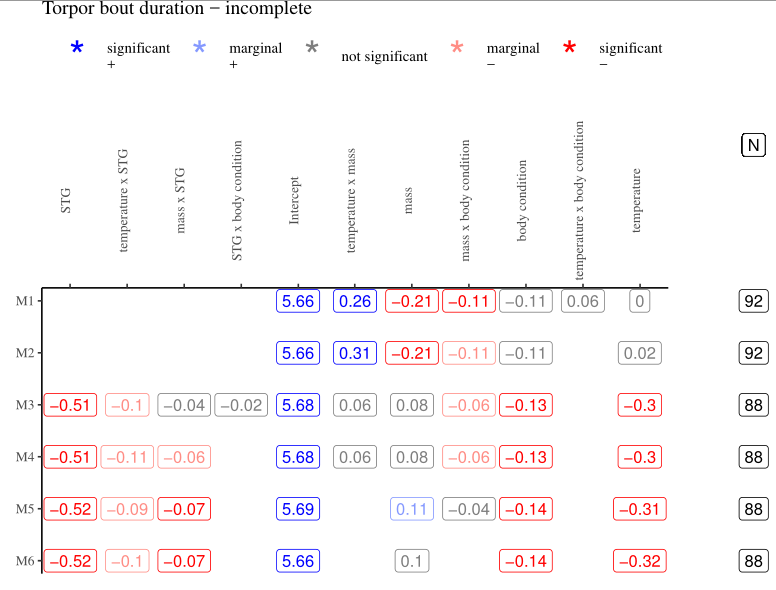

***Figure S23.*** *Graphical illustration of the model simplification process for variables that explained variation in incomplete torpor bout duration.* M1 represents the full model. N shows the variation in sample size due to considering the variables of pre-torpor and torpor STG.

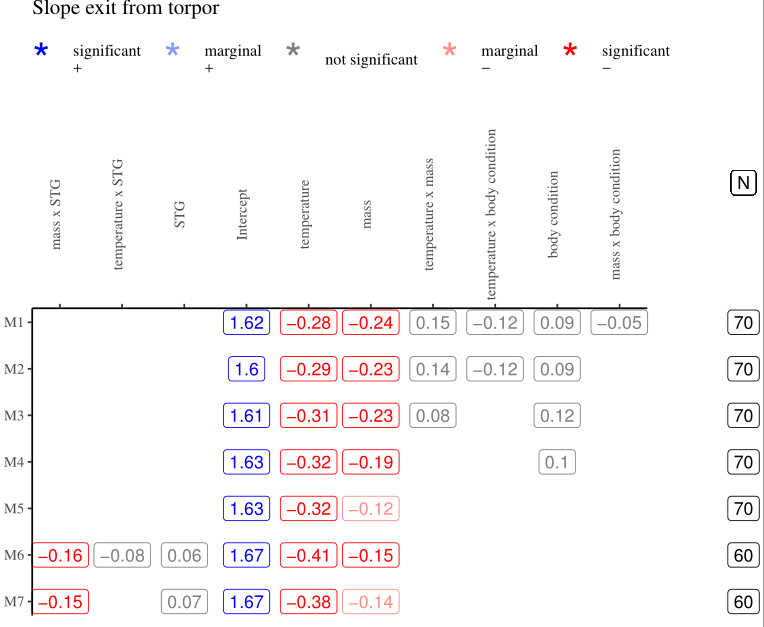

***Figure S24.*** *Graphical illustration of the model simplification process for variables that explained variation in rate of exit from torpor.* M1 represents the full model. N shows the variation in sample size due to considering the variables of pre-torpor and torpor STG.

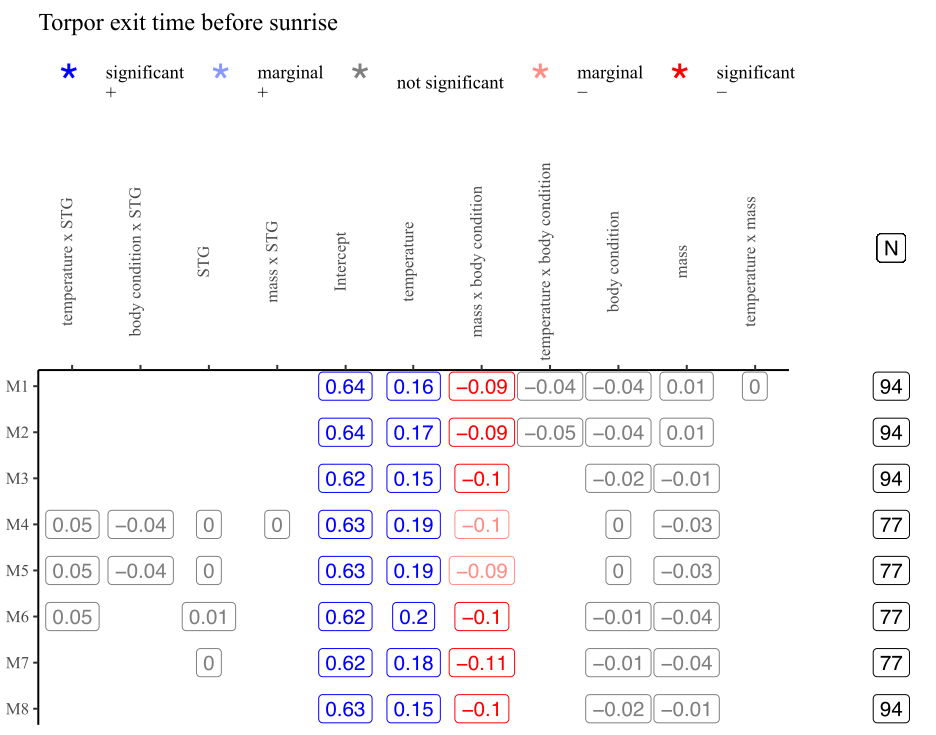

***Figure S25.*** *Graphical illustration of the model simplification process for variables that explained variation in time of torpor initiation.* M1 represents the full model. N shows the variation in sample size due to considering the variables of pre-torpor and torpor STG.

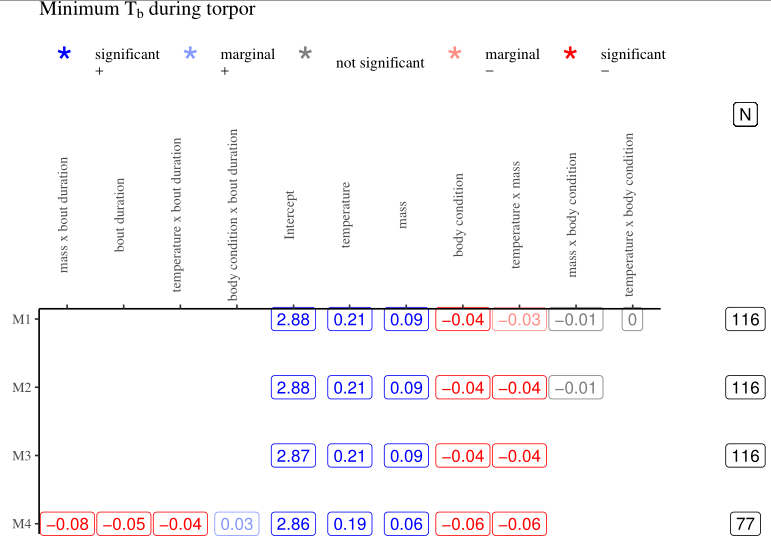

***Figure S26.*** *Graphical illustration of the model simplification process for variables that explained variation in minimum body temperature during torpor (T_b_ min).* M1 represents the full model. N shows the variation in sample size due to considering the variables of pre-torpor and torpor STG.

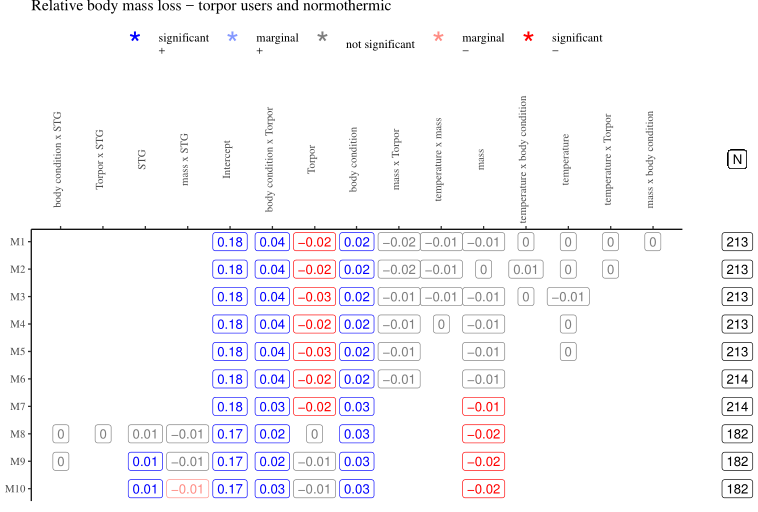

***Figure S27.*** *Graphical illustration of the model simplification process for variables that explained variation in relative mass loss for torpid and normothermic birds.* M1 represents the full model. N shows the variation in sample size due to considering the variable of STG over the entire night. Torpor retains a significant negative effect until STG is considered (compare coefficients in M7-M9).

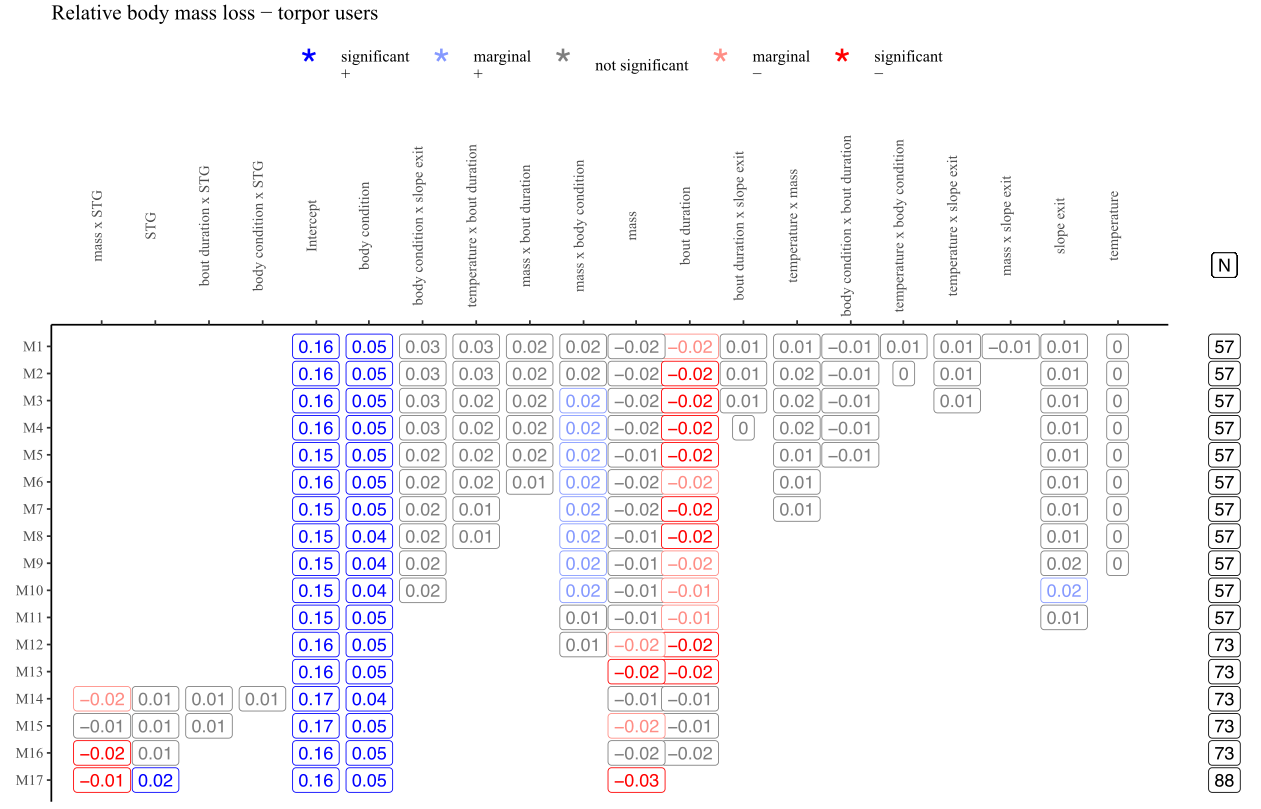

***Figure S28.*** *Graphical illustration of the model simplification process for variables that explained variation in relative mass loss for torpid birds.* M1 represents the full model. N shows the variation in sample size due to considering the variables of STG over the entire night and torpor bout duration. Torpor bout duration retains a significant negative effect until STG is considered (compare coefficients in M13-M17).

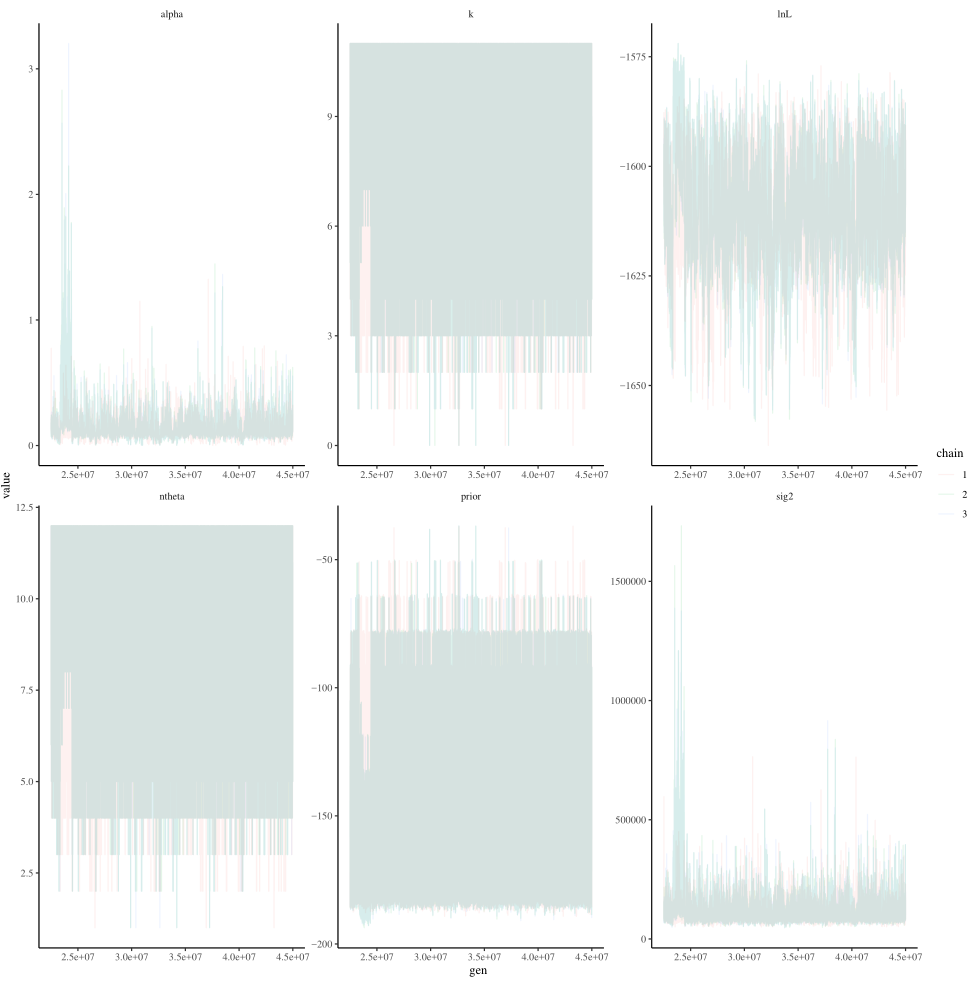

***Figure S29.*** *Traceplots of bayou candidate model parameters.*

**Supplementary Tables**

***Table S1.*** *Coefficient table for models of torpor usage.* Posterior medians in bold are at least marginally significant. *indicates terms that were eliminated in model selection, ᵻ indicates terms that were obtained from the reduced dataset and x indicates interaction. Body mass is initial body mass, T_a_ is median T_a_, and STG is the sustained temperature gradient during the pre-torpor phase (Fig. S14A).

| **Parameter** | **Median** | **95% CI** | | **90% CI** | |
| --- | --- | --- | --- | --- | --- |
|  |  | **Lower** | **Upper** | **Lower** | **Upper** |
| ᵻBody mass | **-1.180** | -2.260 | -0.446 | -2.060 | -0.610 |
| ᵻT_a_ | **-2.380** | -3.640 | -1.410 | -3.430 | -1.630 |
| ᵻBody condition | **0.556** | -0.080 | 1.140 | 0.021 | 1.010 |
| ᵻpre-torpor STG | **-0.992** | -1.990 | -0.224 | -1.800 | -0.398 |
| ᵻBody mass x T_a_ | **0.842** | 0.198 | 1.500 | 0.295 | 1.340 |
| *Body mass x body condition | -0.138 | -0.537 | 0.182 | -0.475 | 0.115 |
| *ᵻBody mass x pre-torpor STG | 0.497 | -0.555 | 1.340 | -0.375 | 1.160 |
| *T_a_ x body condition | 0.124 | -0.539 | 0.679 | -0.498 | 0.554 |
| *ᵻT_a_ x pre-torpor STG | 0.692 | -0.148 | 1.420 | -0.021 | 1.260 |
| *ᵻBody condition x pre-torpor STG | -0.048 | -0.716 | 0.485 | -0.603 | 0.369 |

***Table S2.*** *Coefficient table for models of torpor duration.* Posterior medians in bold are at least marginally significant. *indicates terms that were eliminated in model selection, ᵻ indicates terms that were obtained from the reduced dataset and x indicates interaction. Body mass is initial body mass, T_a_ is median T_a_, and STG is the sustained temperature gradient during the pre-torpor and torpor phases (Fig. S14B).

| **TBD** | **Parameter** | **Median** | **95% CI** | | **90% CI** | |
| --- | --- | --- | --- | --- | --- | --- |
|  |  |  | **Lower** | **Upper** | **Lower** | **Upper** |
| TBD | ᵻBody mass | 0.150 | -0.057 | 0.320 | -0.019 | 0.278 |
| short | ᵻT_a_ | **-0.308** | -0.471 | -0.159 | -0.442 | -0.195 |
| (N = 78) | ᵻBody condition | **-0.264** | -0.401 | -0.155 | -0.378 | -0.179 |
|  | ᵻSTG | **-0.590** | -0.701 | -0.495 | -0.680 | -0.515 |
|  | Body mass x T_a_ | **0.225** | 0.038 | 0.398 | 0.066 | 0.358 |
|  | *Body mass x Body condition | -0.108 | -0.279 | 0.033 | -0.250 | 0.004 |
|  | ᵻBody mass x STG | **-0.114** | -0.234 | -0.017 | -0.211 | -0.037 |
|  | ᵻT_a_ x Body condition | **-0.552** | -0.742 | -0.389 | -0.709 | -0.425 |
|  | *ᵻT_a_ x STG | -0.080 | -0.258 | 0.058 | -0.230 | 0.028 |
|  | *ᵻSTG x Body condition | 0.026 | -0.106 | 0.134 | -0.088 | 0.11 |
| TBD | ᵻBody mass | 0.015 | -0.131 | 0.117 | -0.102 | 0.096 |
| complete | ᵻT_a_ | **-0.247** | -0.401 | -0.107 | -0.377 | -0.142 |
| (N = 63) | *Body condition | -0.085 | -0.291 | 0.085 | -0.257 | 0.046 |
|  | ᵻSTG | **-0.506** | -0.615 | -0.407 | -0.599 | -0.430 |
|  | ᵻBody mass x T_a_ | **0.171** | 0.027 | 0.302 | 0.049 | 0.274 |
|  | *Body mass x Body condition | -0.096 | -0.262 | 0.039 | -0.232 | 0.008 |
|  | *ᵻBody mass x STG | -0.048 | -0.169 | 0.052 | -0.148 | 0.028 |
|  | *T_a_ x Body condition | -0.123 | -0.658 | 0.325 | -0.562 | 0.219 |
|  | ᵻT_a_ x STG | **-0.179** | -0.369 | -0.018 | -0.339 | -0.057 |
| TBD | ᵻBody mass | 0.098 | -0.030 | 0.202 | -0.007 | 0.180 |
| incomplete | ᵻT_a_ | **-0.318** | -0.447 | -0.203 | -0.428 | -0.231 |
| (N = 93) | ᵻBody condition | **-0.136** | -0.240 | -0.043 | -0.222 | -0.064 |
|  | ᵻSTG | **-0.522** | -0.617 | -0.445 | -0.600 | -0.463 |
|  | *ᵻBody mass x T_a_ | 0.061 | -0.056 | 0.151 | -0.039 | 0.132 |
|  | *ᵻBody mass x Body condition | -0.04 | -0.114 | 0.021 | -0.103 | 0.007 |
|  | ᵻBody mass x STG | **-0.074** | -0.153 | -0.007 | -0.14 | -0.021 |
|  | *T_a_ x Body condition | 0.063 | -0.151 | 0.244 | -0.120 | 0.203 |
|  | T_a_ x STG | **-0.103** | -0.237 | 0.009 | -0.211 | -0.017 |
|  | *ᵻSTG x Body condition | -0.02 | -0.127 | 0.065 | -0.109 | 0.046 |

***Table S3.*** *Coefficient table for models of rate of exit from torpor, time of torpor exit and minimum T_b_ during torpor.* Posterior medians in bold are at least marginally significant. *indicates terms that were eliminated in model selection, ᵻ indicates terms that were obtained from the reduced dataset and x indicates interaction. Body mass is initial body mass, T_a_ is median T_a_, and STG is the sustained temperature gradient during the pre-torpor and torpor phases (Fig. S14B).

| **Torpor characteristic** | **Parameter** | **Median** | **95% CI** | | **90% CI** | |
| --- | --- | --- | --- | --- | --- | --- |
|  |  |  | **Lower** | **Upper** | **Lower** | **Upper** |
| Rate | ᵻBody mass | **-0.145** | -0.310 | 0.003 | -0.284 | -0.032 |
| of | ᵻT_a_ | **-0.381** | -0.549 | -0.238 | -0.522 | -0.271 |
| Exit | *Body condition | 0.097 | -0.057 | 0.227 | -0.033 | 0.198 |
|  | ᵻSTG | 0.072 | -0.082 | 0.192 | -0.056 | 0.168 |
|  | *Body mass x T_a_ | 0.078 | -0.061 | 0.196 | -0.038 | 0.168 |
|  | *Body mass x Body condition | -0.048 | -0.172 | 0.057 | -0.150 | 0.033 |
|  | ᵻBody mass x STG | **-0.153** | -0.304 | -0.028 | -0.278 | -0.056 |
|  | *T_a_ x Body condition | -0.116 | -0.343 | 0.079 | -0.304 | 0.033 |
|  | *ᵻT_a_ x STG | -0.082 | -0.316 | 0.108 | -0.276 | 0.071 |
| Time | ᵻBody mass | -0.009 | -0.173 | 0.122 | -0.147 | 0.096 |
| of | ᵻT_a_ | **0.149** | 0.018 | 0.260 | 0.042 | 0.235 |
| exit | ᵻBody condition | -0.023 | -0.139 | 0.073 | -0.121 | 0.052 |
| initiation | *ᵻSTG | 0.003 | -0.106 | 0.092 | -0.090 | 0.073 |
| (h before | *Body mass x T_a_ | -0.005 | -0.159 | 0.122 | -0.134 | 0.094 |
| sunrise) | ᵻBody mass x Body condition | **-0.098** | -0.182 | -0.026 | -0.169 | -0.041 |
|  | *ᵻBody mass x STG | 0.001 | -0.152 | 0.125 | -0.126 | 0.096 |
|  | *T_a_ x Body condition | -0.011 | -0.159 | 0.148 | -0.132 | 0.116 |
|  | *ᵻT_a_ x STG | 0.05 | -0.136 | 0.204 | -0.104 | 0.170 |
|  | *ᵻSTG x Body condition | -0.04 | -0.145 | 0.049 | -0.128 | 0.029 |
| Minimum | ᵻBody mass | **0.063** | 0.014 | 0.108 | 0.021 | 0.097 |
| T_b_ | ᵻT_a_ | **0.191** | 0.147 | 0.228 | 0.154 | 0.220 |
|  | ᵻBody condition | **-0.063** | -0.103 | -0.032 | -0.097 | -0.039 |
|  | ᵻTBD (short) | **-0.054** | -0.086 | -0.080 | -0.080 | -0.034 |
|  | ᵻBody mass x T_a_ | **-0.065** | -0.112 | -0.028 | -0.103 | -0.036 |
|  | *Body mass x Body condition | -0.013 | -0.039 | 0.01 | -0.034 | 0.005 |
|  | ᵻBody mass x TBD | **-0.083** | -0.125 | -0.049 | -0.118 | -0.057 |
|  | *T_a_ x Body condition | -0.004 | -0.045 | 0.031 | -0.038 | 0.023 |
|  | ᵻT_a_ x TBD | **-0.035** | -0.069 | -0.007 | -0.064 | -0.013 |
|  | ᵻTBD x Body condition | **0.033** | -0.002 | 0.065 | 0.003 | 0.058 |

***Table S4.*** *Coefficient table for models of relative body mass loss.* Posterior medians in bold are at least marginally significant. *indicates terms that were eliminated in model selection, ᵻ indicates terms that were obtained from the reduced dataset and x indicates interaction. Body mass is initial body mass, T_a_ is median T_a_ and STG is the sustained temperature gradient over the whole night (Fig. S14C). TBD is short TBD.

| **Data set** | **Parameter** | **Median** | **95% CI** | | **90% CI** | |
| --- | --- | --- | --- | --- | --- | --- |
|  |  |  | **Lower** | **Upper** | **Lower** | **Upper** |
| All birds | ᵻBody mass | **-0.020** | -0.037 | -0.006 | -0.034 | -0.009 |
|  | *T_a_ | -0.005 | -0.018 | 0.005 | -0.016 | 0.003 |
|  | ᵻBody condition | **0.029** | 0.014 | 0.040 | 0.016 | 0.038 |
|  | ᵻSTG | **0.013** | 0.001 | 0.022 | 0.003 | 0.020 |
|  | *Body mass x T_a_ | -0.003 | -0.016 | 0.007 | -0.014 | 0.004 |
|  | *Body mass x Body condition | 0.001 | -0.009 | 0.007 | -0.007 | 0.005 |
|  | ᵻBody mass x STG | **-0.008** | -0.018 | 0.000 | -0.016 | -0.001 |
|  | *ᵻBody condition x STG | -0.003 | -0.016 | 0.008 | -0.015 | 0.005 |
|  | *T_a_ x Body condition | 0.005 | -0.009 | 0.016 | -0.007 | 0.014 |
|  | ᵻTorpor use | -0.006 | -0.026 | 0.012 | -0.023 | 0.008 |
|  | *Torpor use x Body mass | -0.009 | -0.028 | 0.009 | -0.025 | 0.004 |
|  | Torpor use x Body condition | **0.026** | 0.008 | 0.040 | 0.011 | 0.037 |
|  | *Torpor use x T_a_ | 0.002 | -0.023 | 0.015 | -0.019 | 0.011 |
|  | *ᵻTorpor use x STG | -0.002 | -0.024 | 0.016 | -0.020 | 0.012 |
| Torpid | ᵻBody mass | **-0.025** | -0.046 | -0.009 | -0.043 | -0.013 |
| birds | *T_a_ | -0.001 | -0.030 | 0.024 | -0.025 | 0.018 |
|  | ᵻBody condition | **0.053** | 0.036 | 0.068 | 0.039 | 0.065 |
|  | ᵻSTG | **0.017** | 0.003 | 0.029 | 0.006 | 0.026 |
|  | *ᵻTBD | -0.016 | -0.040 | 0.004 | -0.036 | 0.000 |
|  | *Slope torpor exit | 0.013 | -0.005 | 0.029 | -0.002 | 0.025 |
|  | *Body mass x T_a_ | 0.010 | -0.017 | 0.032 | -0.012 | 0.027 |
|  | *Body mass x Body condition | 0.008 | -0.009 | 0.023 | -0.007 | 0.020 |
|  | ᵻBody mass x STG | **-0.012** | -0.025 | -0.002 | -0.023 | -0.004 |
|  | *Body mass x TBD | 0.012 | -0.013 | 0.031 | -0.009 | 0.027 |
|  | *Body mass x slope torpor exit | -0.008 | -0.038 | 0.017 | -0.033 | 0.012 |
|  | *T_a_ x Body condition | 0.005 | -0.068 | 0.063 | -0.056 | 0.051 |
|  | *T_a_ x TBD | 0.012 | -0.012 | 0.030 | -0.008 | 0.026 |
|  | *T_a_ x slope torpor exit | 0.008 | -0.029 | 0.040 | -0.022 | 0.033 |
|  | *ᵻBody condition x STG | 0.009 | -0.013 | 0.028 | -0.009 | 0.024 |
|  | *Body condition x TBD | -0.009 | -0.033 | 0.010 | -0.030 | 0.006 |
|  | *Body condition x slope torpor exit | 0.021 | -0.010 | 0.047 | -0.004 | 0.041 |
|  | *ᵻSTG x TBD | 0.010 | -0.006 | 0.024 | -0.003 | 0.021 |
|  | *TBD x slope torpor exit | 0.004 | -0.026 | 0.029 | -0.020 | 0.023 |

***Table S5.*** *Variance partitioning from final PGLMM regression models of each torpor trait.*

| **Response variable** | **Proportion of total variance attributable to phylogenetic random effect** | | | **Proportion of total variance attributable to intraspecific variation** | | |
| --- | --- | --- | --- | --- | --- | --- |
|  | Median | Standard error | 95% credible interval | Median | Standard error | 95% credible interval |
| Torpor usage | 0.29 | 0.32 | 0.01 – 0.99 | 0.71 | 0.32 | 0.01-1.0 |
| TBD (short) | 0.01 | 0.02 | 0.01 - 0.08 | 0.33 | 0.23 | 0.01-0.76 |
| TBD (complete) | 0.01 | 0.02 | 0.01 – 0.06 | 0.17 | 0.18 | 0.01-0.65 |
| TBD (incomplete) | 0.01 | 0.01 | 0.01 – 0.02 | 0.1 | 0.11 | 0.01-0.41 |
| Rate of exit | 0.01 | 0.02 | 0.01 – 0.06 | 0.23 | 0.17 | 0.01-0.61 |
| Time of arousal | 0.02 | 0.02 | 0.01 – 0.07 | 0.22 | 0.18 | 0.01-0.61 |
| Minimum T_b_ | 0.01 | 0.02 | 0.01 – 0.06 | 0.21 | 0.19 | 0.01-0.63 |
| Relative mass loss  (all birds) | 0.01 | 0.01 | 0.01 – 0.03 | 0.20 | 0.11 | 0.01-0.44 |
| Relative mass loss  (torpor users) | 0.01 | 0.01 | 0.01 – 0.03 | 0.13 | 0.12 | 0.01-0.41 |

***Table S6.*** *Phylogenetic signal of species-level torpor trait variation that accounts for variation in ecology.* For each response variable, we selected the final PGLMM, extracted the phylogenetic species-level random effect level, and calculated its phylogenetic signal. Pagel’s **λ** was estimated with a Bayesian sample in motmot. Blomberg’s K was calculated with phytools. P-values come from 1,000 iterations of bootstrapping. Errors supplied for alternative parameterization of Blomberg’s K come from posterior estimates of the standard error of the phylogenetic random intercept term.

| **Response variable** | **Pagel’s λ** | | **Blomberg’s K** | | **Blomberg’s K with errors** | |
| --- | --- | --- | --- | --- | --- | --- |
|  | Median | 95% credible interval | K | P | K | P |
| Torpor usage | 0.96 | 0.78 – 0.99 | 2.25 | 0.001 | 1.08 | 0.001 |
| TBD (short) | 0.72 | 0.25 – 0.99 | 1.13 | 0.001 | 1.19 | 0.001 |
| TBD (complete) | 0.83 | 0.49 – 0.99 | 1.64 | 0.001 | 1.25 | 0.001 |
| TBD (incomplete) | 0.80 | 0.36 – 0.99 | 1.26 | 0.001 | 1.14 | 0.001 |
| Rate of exit | 0.67 | 0.13 – 0.99 | 0.86 | 0.047 | 1.21 | 0.001 |
| Time of arousal | 0.86 | 0.50 – 0.99 | 1.45 | 0.001 | 1.15 | 0.001 |
| Minimum T_b_ | 0.48 | 0.03 - 0.96 | 0.80 | 0.040 | 1.23 | 0.001 |
| Relative mass loss (torpor vs. normothermic) | 0.62 | 0.18 – 0.99 | 0.95 | 0.001 | 1.14 | 0.001 |
| Relative mass loss  (torpor users) | 0.59 | 0.16 – 0.97 | 1.06 | 0.002 | 1.08 | 0.001 |

***Table S7****. Summary statistics of bayou candidate model indicate convergence.* Three chains ran for 45,000,000 iterations and a burn-in of 90%. SD shows standard deviation, HPD shows region of 95% highest posterior density and ESS shows effective sample size.

| **Term** | **Mean** | **SD** | **HPD2.5%** | **HPD97.5%** | **ESS** |
| --- | --- | --- | --- | --- | --- |
| lnL | -1608.684 | 9.772 | -1628.784 | -1589.182 | 750.636 |
| Prior | -132.870 | 24.848 | -184.329 | -90.958 | 8295.952 |
| Alpha | 0.133 | 0.054 | 0.038 | 0.235 | 2276.037 |
| Sigma^2^ | 111724.7 | 29138.89 | 66973.68 | 165610.4 | 6074.574 |
| k | 7.096 | 1.870 | 3 | 10 | 8365.612 |
| N_theta_ | 8.096 | 1.870 | 3 | 10 | 8365.612 |
| Root_theta_ | 114.968 | 150.275 | 880.589 | 1417.861 | 682.840 |

***Table S8.*** *An evolutionary model of elevational range reveals multiple, robust shifts in elevation.* Branches of bayou candidate model with locations of elevational shifts that have higher than 10% posterior probabilities (PP) across three chains ran for 45,000,000 iterations and a burn-in of 90%. SE shows standard error. Relative location shows how far down the branch the shift occurs. Descendant taxa denotes taxa that are affected by the shift.

| **Branch number** | **PP** | **Magnitude of theta_2_** | **Naïve SE of theta_2_** | **Relative location** | **Descendant taxa** |
| --- | --- | --- | --- | --- | --- |
| 266 | 0.921 | 2627.543 | 0.224 | 0.658 | Heliantheini |
| 261 | 0.888 | 3249.469 | 0.379 | 0.607 | Lesbiini |
| 137 | 0.719 | 1200.905 | 0.284 | 0.535 | *Heliodoxa sp. and Urochroa sp.* |
| 401 | 0.561 | 4228.047 | 1.212 | 0.430 | *Patagona gigas* |
| 136 | 0.154 | 1016.913 | 0.767 | 0.528 | *Heliodoxa sp.* |

***Table S9.*** *A constrained null model of elevational range fails to reveal the same shifts as the candidate model.* Branches of bayou null model with constrained shift numbers showing locations of elevational shifts that have higher than 10% posterior probabilities (PP) across one chain ran for 10,000,000 iterations and a burn-in of 50%. SE shows standard error. Relative location shows how far down the branch the shift occurs. Descendant taxa denotes taxa that are affected by the shift.

| **Branch number** | **PP** | **Magnitude of theta_2_** | **Naïve SE of theta_2_** | **Relative location** | **Descendant taxa** |
| --- | --- | --- | --- | --- | --- |
| 261 | 0.444 | 4316.819 | 2.033 | 0.529 | Lesbiini |
| 266 | 0.299 | 2669.864 | 1.619 | 0.513 | Heliantheini |

**References**

Baldwin, J.W., Garcia-Porta, J. & Botero, C.A. (2022). Phenotypic responses to climate change are significantly dampened in big-brained birds. *Ecol. Lett.*, 25, 939–947.

Barrow, L.N., McNew, S.M., Mitchell, N., Galen, S.C., Lutz, H.L., Skeen, H., *et al.* (2019). Deeply conserved susceptibility in a multi-host, multi-parasite system. *Ecol. Lett.*, 22, 987–998.

Bolstad, G.H., Hansen, T.F., Pélabon, C., Falahati-Anbaran, M., Pérez-Barrales, R. & Armbruster, W.S. (2014). Genetic constraints predict evolutionary divergence in Dalechampia blossoms. *Philos. Trans. R. Soc. B Biol. Sci.*, 369.

Brigham, R.M. (1992). Daily torpor in a free-ranging goatsucker, the common poorwill (Phalaeonoptilus nuttallii). *Physiol. Zool.*, 65, 457–472.

Brigham, R.M., Körtner, G., Maddocks, T.A. & Geiser, F. (2000). Seasonal use of torpor by free-ranging Australian owlet-nightjars (Aegotheles cristatus). *Physiol. Biochem. Zool.*, 73, 613–620.

Bürkner, P.C. (2017). brms: An R package for Bayesian multilevel models using Stan. *J. Stat. Softw.*, 80.

Bürkner, P.C. (2018). Advanced Bayesian multilevel modeling with the R package brms. *R J.*, 10, 395–411.

Calder, W.A. (1994). When do hummingbirds use torpor in nature? *Physiol. Zool.*, 67, 1051–1076.

Carpenter, B., Gelman, A., Hoffman, M.D., Lee, D., Goodrich, B., Betancourt, M., *et al.* (2017). Stan: A probabilistic programming language. *J. Stat. Softw.*, 76, 1–32.

Chang, W., Cheng, J., Allaire, J.J., Sievert, C., Schloerke, B., Xie, Y., *et al.* (2021). shiny: Web Application Framework for R.

Dormann, C.F., Elith, J., Bacher, S., Buchmann, C., Carl, G., Carré, G., *et al.* (2013). Collinearity: A review of methods to deal with it and a simulation study evaluating their performance. *Ecography (Cop.).*, 36, 27–46.

Fristoe, T.S., Burger, J.R., Balk, M.A., Khaliq, I., Hof, C. & Brown, J.H. (2015). Metabolic heat production and thermal conductance are mass-independent adaptations to thermal environment in birds and mammals. *Proc. Natl. Acad. Sci. U. S. A.*, 112, 15934–15939.

Garland, T. & Ives, A.R. (2000). Using the past to predict the present: Confidence intervals for regression equations in phylogenetic comparative methods. *Am. Nat.*, 155, 346–364.

Geiser, F. (2004). Metabolic rate and body temperature reduction during hibernation and daily torpor. *Annu. Rev. Physiol.*, 66, 239–274.

Gilbert, C., Blanc, S., Le Maho, Y. & Ancel, A. (2008). Energy saving processes in huddling emperor penguins: From experiments to theory. *J. Exp. Biol.*, 211, 1–8.

Gilbert, C., McCafferty, D., Le Maho, Y., Martrette, J.M., Giroud, S., Blanc, S., *et al.* (2010). One for all and all for one: The energetic benefits of huddling in endotherms. *Biol. Rev.*, 85, 545–569.

Graham, C.H., Parra, J.L., Rahbek, C. & McGuire, J.A. (2009). Phylogenetic structure in tropical hummingbird communities. *Proc. Natl. Acad. Sci. U. S. A.*, 106, 19673–19678.

Hackett, S.J., Kimball, R.T., Reddy, S., Bowie, R.C.K., Braun, E.L., Braun, M.J., *et al.* (2008). A phylogenomic study of birds reveals their evolutionary history. *Science (80-. ).*, 320, 1763–1768.

Hansen, T.F., Bolstad, G.H. & Tsuboi, M. (2021). Analyzing Disparity and Rates of Morphological Evolution with Model-Based Phylogenetic Comparative Methods. *Syst. Biol.*, 0, 1–19.

Jetz, W., Thomas, G.H., Joy, J.B., Hartmann, K. & Mooers, A.O. (2012). The global diversity of birds in space and time. *Nature*, 491, 444–448.

Krementz, D.G. & Pendleton, G.W. (1990). Fat Scoring: Sources of Variability. *Condor*, 92, 500–507.

Londoño, G.A., Chappell, M.A., Jankowski, J.E. & Robinson, S.K. (2017). Do thermoregulatory costs limit altitude distributions of Andean forest birds? *Funct. Ecol.*, 31, 204–215.

McGuire, J.A., Witt, C.C., Remsen, J. V., Corl, A., Rabosky, D.L., Altshuler, D.L., *et al.* (2014). Molecular phylogenetics and the diversification of hummingbirds. *Curr. Biol.*, 24, 910–916.

Mckechnie, A.E. & Lovegrove, B.G. (2002). Avian Facultative Hypothermic Responses: A Review. *The Condor2*, 104, 705–724.

Paterno, G.B., Penone, C. & Werner, G.D.A. (2018). sensiPhy: An r-package for sensitivity analysis in phylogenetic comparative methods. *Methods Ecol. Evol.*, 9, 1461–1467.

Pennell, M.W., Eastman, J.M., Slater, G.J., Brown, J.W., Uyeda, J.C., Fitzjohn, R.G., *et al.* (2014). Geiger v2.0: An expanded suite of methods for fitting macroevolutionary models to phylogenetic trees. *Bioinformatics*, 30, 2216–2218.

Porter, W.P. (1969). Thermal Radiation in Metabolic Chambers. *Science (80-. ).*, 166, 115–117.

Quintero, I. & Jetz, W. (2018). Global elevational diversity and diversification of birds. *Nature*, 555, 246–250.

R Development Core Team. (2012). R: A language and environment for statistical computing. R Foundation for Statistical Computing.

Revell, L.J. (2012). phytools: An R package for phylogenetic comparative biology (and other things). *Methods Ecol. Evol.*, 3, 217–223.

Ricklefs, R.E. & Williams, J.B. (2003). Metabolic responses of shorebird chicks to cold stress: Hysteresis of cooling and warming phases. *J. Exp. Biol.*, 206, 2883–2893.

Rising, J.D. & Somers, K.M. (1989). The Measurement of Overall Body Size in Birds. *Auk*, 106, 666–674.

Ruf, T. & Geiser, F. (2015). Daily torpor and hibernation in birds and mammals. *Biol. Rev.*, 90, 891–926.

Schleucher, E. (2004). Torpor in birds: Taxonomy, energetics, and ecology. *Physiol. Biochem. Zool.*, 77, 942–949.

Senar, J.C. & Pascual, J. (1997). Keel and tarsus length may provide a good predictor of avian body size. *Ardea*, 85, 269–274.

Shankar, A., Cisneros, I.N.H., Thompson, S., Graham, C.H. & Powers, D.R. (2022). A heterothermic spectrum in hummingbirds. *J. Exp. Biol.*, 225, 1–10.

Stiles, F.G. (2004). Phylogenetic Constraints Upon Morphological and Ecological Adaptation in Hummingbirds ( Trochilidae ): Why Are There No Hermits in the Paramo ? *Ornitol. Netropical*, 15, 191–198.

Thomas, G.H. & Freckleton, R.P. (2012). MOTMOT: Models of trait macroevolution on trees. *Methods Ecol. Evol.*, 3, 145–151.

Tobias, J.A., Sheard, C., Pigot, A.L., Devenish, A.J.M., Yang, J., Neate-Clegg, M.H.C., *et al.* (2022). AVONET: morphological, ecological and geographical data for all birds. *Ecol. Lett.*, 25, 581–597.

Tomlinson, S., Withers, P.C. & Cooper, C. (2007). Hypothermia versus torpor in response to cold stress in the native Australian mouse Pseudomys hermannsburgensis and the introduced house mouse Mus musculus. *Comp. Biochem. Physiol. - A Mol. Integr. Physiol.*, 148, 645–650.

Tung Ho, L.S. & Ané, C. (2014). A linear-time algorithm for gaussian and non-gaussian trait evolution models. *Syst. Biol.*, 63, 397–408.

Uyeda, J.C. & Harmon, L.J. (2014). A novel Bayesian method for inferring and interpreting the dynamics of adaptive landscapes from phylogenetic comparative data. *Syst. Biol.*, 63, 902–918.

Uyeda, J.C., Pennell, M.W., Miller, E.T., Maia, R. & McClain, C.R. (2017). The evolution of energetic scaling across the vertebrate tree of life. *Am. Nat.*, 190, 185–199.

de Villemereuil, P. & Nakagawa, S. (2014). Modern phylogenetic comparative methods and their application in evolutionary biology. In: *Modern Phylogenetic Comparative Methods and Their Application in Evolutionary Biology*. pp. 287–303.

Wang, L.C.H. (1989). Ecological, physiological, and biochemical aspects of torpor in mammals and birds. In: *Animal Adaptation to cold*. Springer Berlin Heidelberg, Berlin, pp. 361–401.

Wilman, H., J., B., J., S., C., de L.R., M., R. & W, J. (2014). EltonTraits 1 . 0 : Species-level foraging attributes of the world ’ s birds and mammals. *Ecology*, 95, 2027.

Wolf, B.O., Mckechnie, A.E., Schmitt, C.J., Czenze, Z.J., Johnson, A.B., Witt, C.C., *et al.* (2020). Extreme and variable torpor among high- elevation Andean hummingbird species. *Biol. Lett.*, 5–9.

Wolf, B.O. & Walsberg, G.E. (2000). The role of the plumage in heat transfer processes of birds. *Am. Zool.*, 40, 575–584.
